# Nucleoporins shape germ granule architecture and balance small RNA silencing pathways

**DOI:** 10.1101/2025.01.23.634177

**Authors:** Kun Shi, Ying Zhang, Zhenzhen Du, Symonne C Liu, Xinyu Fan, Heng-Chi Lee, Donglei Zhang

## Abstract

Animals have evolved distinct small RNA pathways, including piRNA and siRNA, to silence invasive and selfish nucleic acids. piRNA pathway factors are concentrated in perinuclear germ granules that frequently associate with nuclear pore complexes (NPCs). However, the factors mediating germ granule-NPC association and the functional relevance of such association remain unknown. Here we show that the conserved nucleoporins NPP-14 (NUP-214) and NPP-24 (NUP-88), components of the cytoplasmic filaments of NPC, play critical roles in anchoring germ granule to NPC and in attenuating piRNA silencing In *C. elegans*. Proximity labeling experiments further identified EPS-1 (enhanced piRNA silencing) as a key germ granule factor contributing to germ granule-NPC interaction. In *npp-14*, *npp-24,* or *eps-1* mutant animals, we observed fewer but enlarged, unorganized germ granules, accompanied by the over-amplification of secondary small RNAs at piRNA targeting sites. Nonetheless, we found this enhancement of piRNA silencing comes at the cost of dampened RNAi efficiency and RNAi inheritance. Together, our studies uncovered factors contributing to germ granule-NPC association and underscored the importance of spatial organization of germ granules in balancing small RNA silencing pathways.

## INTRODUCTION

Small RNA pathways play crucial roles in genome defense by silencing invasive nucleic acids such as transposons and viruses^1^. Among these pathways, PIWI-interacting RNAs (piRNAs) and small interfering RNAs (siRNAs) have been extensively studied for their involvement in safeguarding genome integrity^2^. In the nematode *Caenorhabditis elegans (C. elegans)*, both piRNA and siRNA pathways operate in the defense against transposable elements^3^ and share common downstream components^4,5^, suggesting potential crosstalk and coordinated regulation between them. Indeed, in PIWI *prg-1* mutant, which lack all piRNAs, the duration of RNAi (siRNA) inheritance is dramatically enhanced^6^. However, whether there is regulatory mechanism controlling the efficacy of piRNAs is not known.

Central to piRNA-mediated gene regulation is the organization of the piRNA pathway factors within specialized cytoplasmic structures known as germ granules^7^. These membrane-less organelles serve as hubs for RNA metabolism and regulation in the germline^8^. Germ granules exhibit a remarkable organization into distinct sub-compartments, including P granules, Z granules, SIMAR foci, and mutator foci, each with specific molecular factors involved in distinct steps of small RNA silencing^9–12^. These observations suggest that distinct germ granule sub-compartments may coordinate to ensure proper initiation and amplification of gene silencing signals.

Germ granules have been observed to be closely associated with nuclear pores in various animals, including zebrafishes, worms, flies, and mice^13–16^. In *C. elegans* germline, germ granules are shown to be the extensions of nuclear pore complexes (NPCs) and serve as major sites for RNA export^17,18^. Despite the association between germ granule and NPCs have been reported for decades, several key questions remain unanswered; what are the factors mediating germ granule and NPC interactions? In addition, what is the functional significance of such interactions in small RNA-mediated silencing? Previous studies examining the contribution of NPC factor nucleoporins in germ granule assembly focus on analyzing the early embryos of *C. elegans*^19,20^. Nonetheless, as several of the key piRNA factors are only enriched in the germ granule in the adult germline but not in the early embryo^21,22^, analyses are needed to examine the germ granule-NPC association in the adult germline in order to investigate their potential roles in small RNA gene silencing.

## RESULTS

### Nucleoporin NPP-14 (NUP-214) and NPP-24 (NUP-88) negatively regulate piRNA silencing

In our previous investigations, we utilized GFP-based piRNA reporters to identify factors that positively regulate piRNA-mediated gene silencing^23,24^. Intriguingly, our previous observations indicated that knockdown of certain nucleoporins (NPPs) may potentiate piRNA silencing^23^. To identify the negative regulators of piRNA silencing, we conducted a targeted RNAi screen encompassing various NPPs to assess their impact on the piRNA reporter silencing at an elevated temperature where reporter silencing is compromised (Fig. S1A and S1B). In this piRNA silencing reporter, the GFP-targeting piRNA effectively silences GFP expression at 20°C^25^, yet such silencing efficacy diminishes at 25°C (Fig. S1A), likely attributable to reduced piRNA expression levels at the elevated temperature^26^. Our screening revealed that RNAi targeting several nucleoporin genes, including *npp-14* and *npp-24,* significantly increased the proportion of animals exhibiting partial or complete silencing of the piRNA reporter at 25°C (Fig. S1C). Notably, RNAi of numerous other nucleoporin genes failed to affect piRNA reporter silencing at 25°C, despite pronounced sterility phenotypes (Fig. S1B and S1C). We further characterized the roles of NPP-14 and NPP-24 since *npp-14* and *npp-24* knockdown animals were the only RNAi-treated animals in our screen that had normal viability but exhibited enhanced piRNA silencing. NPP-14 and NPP-24 are homologues of human NUP214 and NUP88, respectively, which are known to localize at the cytoplasmic filaments of NPC (Fig. 1A)^27^. We further corroborate the roles of *npp-14* and *npp-24* in piRNA silencing using null mutants created by CRISPR-Cas9 (Fig. 1B and 1C). Given that both the *npp-14(dz7)* and the *npp-24(dz12)* null mutants exhibited normal viability with a brood size comparable to that of the wild type (Fig. S1D), it indicates that the overall functionality of NPCs remains intact in the absence of NPP-14 or NPP-24. To ascertain whether the silencing of the GFP piRNA reporter in *npp-14* is contingent upon piRNA activity, we knock-outed the PIWI Argonaute gene *prg-1*, followed by *npp-14*, in the piRNA reporter strain (Fig. 1D). We observed that these *npp-14* mutant animals do not trigger gene silencing of GFP reporter in the absence of *prg-1* (Fig. 1E), indicating that NPP-14 contribute to GFP reporter silencing in a piRNA-dependent manner. Conversely, when *npp-14* was knock-outed prior to *prg-1*, the piRNA reporter remained silenced (Fig. 1D and 1E). It is known that piRNA silencing, once established, can be maintained without piRNAs for generations^28,29^. This finding suggests that once established, the silencing state of the piRNA reporter can be maintained independently of piRNA in this *npp-14* mutant. Collectively, our candidate RNAi screen unveiled NPP-14 and NPP-24 nucleoporins as negative regulators of piRNA silencing.

**Figure 1.**
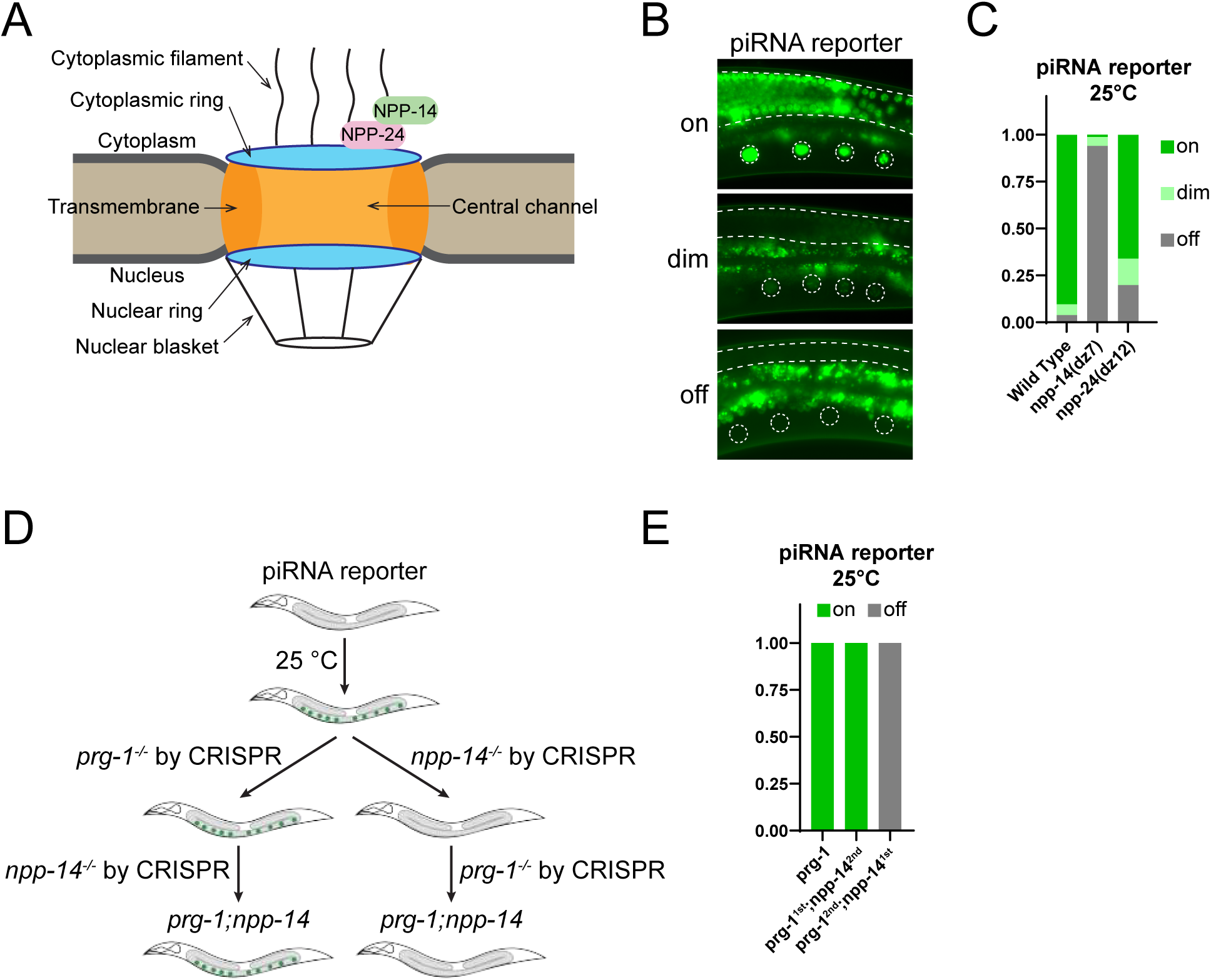
NPP-14 and NPP-24 negatively regulate piRNA silencing. (A) A schematic illustrating the structure of the nuclear pore complex, highlighting NPP-14 (NUP-214 homologue) and NPP-24 (NUP-88 homologue) as nucleoporins located at the cytoplasmic filament. (B) Representative fluorescent micrographs of the piRNA reporter showing different GFP expression levels. In this reporter, the 3’ UTR sequences of *gfp::his-58* are targeted by the endogenous piRNA 21ur-1. Dashed lines and circles denote the germline and maturing oocyte nuclei, respectively. Note that the bright signals observed outside of dashed areas are autofluorescence signals originating from gut granules. (C) Percentage of animals screened displaying the indicated levels of GFP expression levels of the piRNA reporter in the indicated strains grown at 25°C. (D) A schematic illustrating the piRNA reporter assays used to determine whether the initiation or maintenance of reporter silencing in the *npp-14* mutant require PIWI PRG-1. (E) Percentage of animals screened displaying the indicated GFP expression levels of the piRNA reporter in the indicated strains grown at 25°C. *prg-1* and *npp-14* were sequentially knocked out by CRISPR-Cas9 editing, and the order of knockout was indicated.

### NPP-14 and NPP-24 promote the nuclear clustering and tethering of cytoplasmic nucleoporin NPP-9 in the germline

In the *C. elegans* germline, nuclear pores are not randomly distributed but form clusters under germ granules^14^. We monitored the localization of few nucleoporins present in the cytoplasmic filaments of NPC, including RFP::NPP-14, and RFP::NPP-24 and GFP::NPP-9^27^ and confirmed that they all localize as clusters at the nuclear envelope in the germline (Fig. 2A). Consistent with previous report, these NPC clusters were absent in early embryo cells (Fig. S2A)^20^. We also noticed that NPP-14 and NPP-24 are mutually required for each other’s abundance in the germline (Fig. 2A-C and S2B). However, such mutual dependence appears to be specific to germline cells but not in 4-cell embryos (Fig. S2C). In addition, the abundance of other nucleoporins, such as GFP::NPP-9, mCherry::NPP-1, and GFP::NPP-22, were not affected in the germ cells of *npp-14(dz7)* or the *npp-24(dz12)* mutant (Fig. 2A-C, S2D, and S2E).

**Figure 2.**
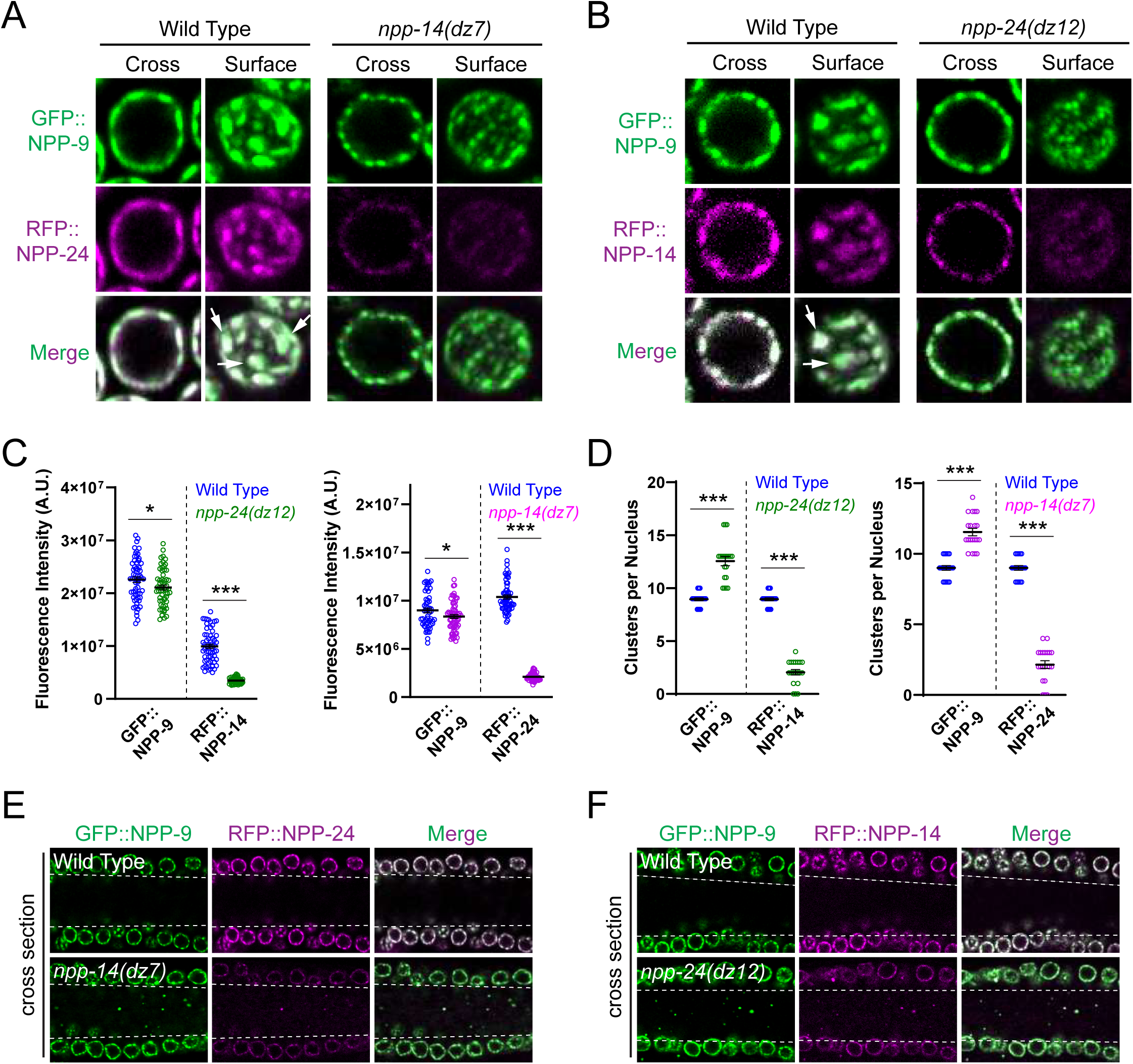
NPP-14 and NPP-24 contribute to NPC clustering and are mutually required for their abundance in the germline (A) Fluorescent micrographs showing the clustering and expression levels of RFP::NPP-24 and GFP::NPP-9 in the indicated strains. The cross and surface sections of germline nuclei from the pachytene region are shown. Arrows indicate NPC clusters. (B) Fluorescent micrographs showing the clustering and expression levels of RFP::NPP-14 and GFP::NPP-9 in the indicated strains. The cross and surface sections of germline nuclei from the pachytene region are shown. Arrows indicate NPC clusters. (C) Mean number of fluorescent intensity of granules in the specified strains. Statistical analysis was conducted using a one-tailed Student’s t-test. n=12-15. *, p<0.05; ***, p<0.001. (D) Mean number of clusters per nucleus in the specified strains. Statistical analysis was conducted using a one-tailed Student’s t-test. n=12-15. *, p<0.05; ***, p<0.001. (E-F) Fluorescent micrographs of the germline cross-section showing the localization of GFP::NPP-9 and (E) RFP::NPP-24 or (F) RFP::NPP-14 in the indicated strains. The area between two dashed lines is germline syncytial cytoplasm.

Interestingly, in the *npp-14(dz7)* or in the *npp-24(dz12)* mutant, we observed an reduction in the size but an increase in the number of GFP::NPP-9 foci at the nuclear envelope (Fig. 2A, 2B, and 2D). Smaller but more NPP-22 foci were also found in *npp-14(dz7)* mutant (Fig. S2F). These results suggest that NPC clustering is compromised in these *npp-14* and in *npp-24* mutants. Furthermore, while most GFP::NPP-9 foci remained associated with the nucleus, a fraction of GFP::NPP-9 foci was detected in the germline syncytial cytoplasm in *npp-14(dz7)* and *npp-24(dz12)* mutants (Fig. 1E and 1F), suggesting that NPP-14 and NPP-24 also promote the tethering of cytoplasmic nucleoporin NPP-9 to the NPCs. In contrast, central channel nucleoporins, such as mCherry::NPP-1 and the transmembrane nucleoporin GFP::NPP-22, remained exclusively associated with the nucleus (Fig. S2G). These results indicate that NPP-14 and NPP-24 contribute to both the NPC clustering as well as the tethering of cytoplasmic nucleoporin NPP-9 to NPC in the germline.

### NPP-14 promotes organization and nuclear tethering of germ granules

As mentioned above, previous studies have demonstrated that germ granules assemble at NPC clusters in the germline^14,17^. In addition, disruption of P granules compromises piRNA silencing^30^. Given our findings of the regulatory roles of NPP-14 and NPP-24 in both piRNA silencing and nuclear pore clustering, we investigated whether NPP-14 and NPP-24 regulate the assembly of germ granules. In wild-type animals, P granule marker mRuby::PGL-1 foci localize on the cytoplasmic side of GFP::NPP-9 foci, consistent with previous reports (Fig. 3A)^14^. While no PGL-1 focus is observed in the syncytial cytoplasm in the wild-type animals, some PGL-1 foci were observed in the syncytial cytoplasm in the *npp-14(dz7)* mutant (Fig. 3B), indicating a weakened association of germ granules with the nucleus. Notably, fewer but larger perinuclear mRuby::PGL-1 foci were observed in the *npp-14(dz7)* or in the *npp-24(dz12)* mutant (Fig. 3A). We also examined other P granule markers, such as mCherry::GLH-1 and GFP::PRG-1, and observed similar results (Fig. S3A). Immunostaining of endogenous PRG-1 also showed a reduced number but enlarged size of PRG-1 foci in the *npp-14(dz7)* mutant (Fig. S3B).

**Figure 3.**
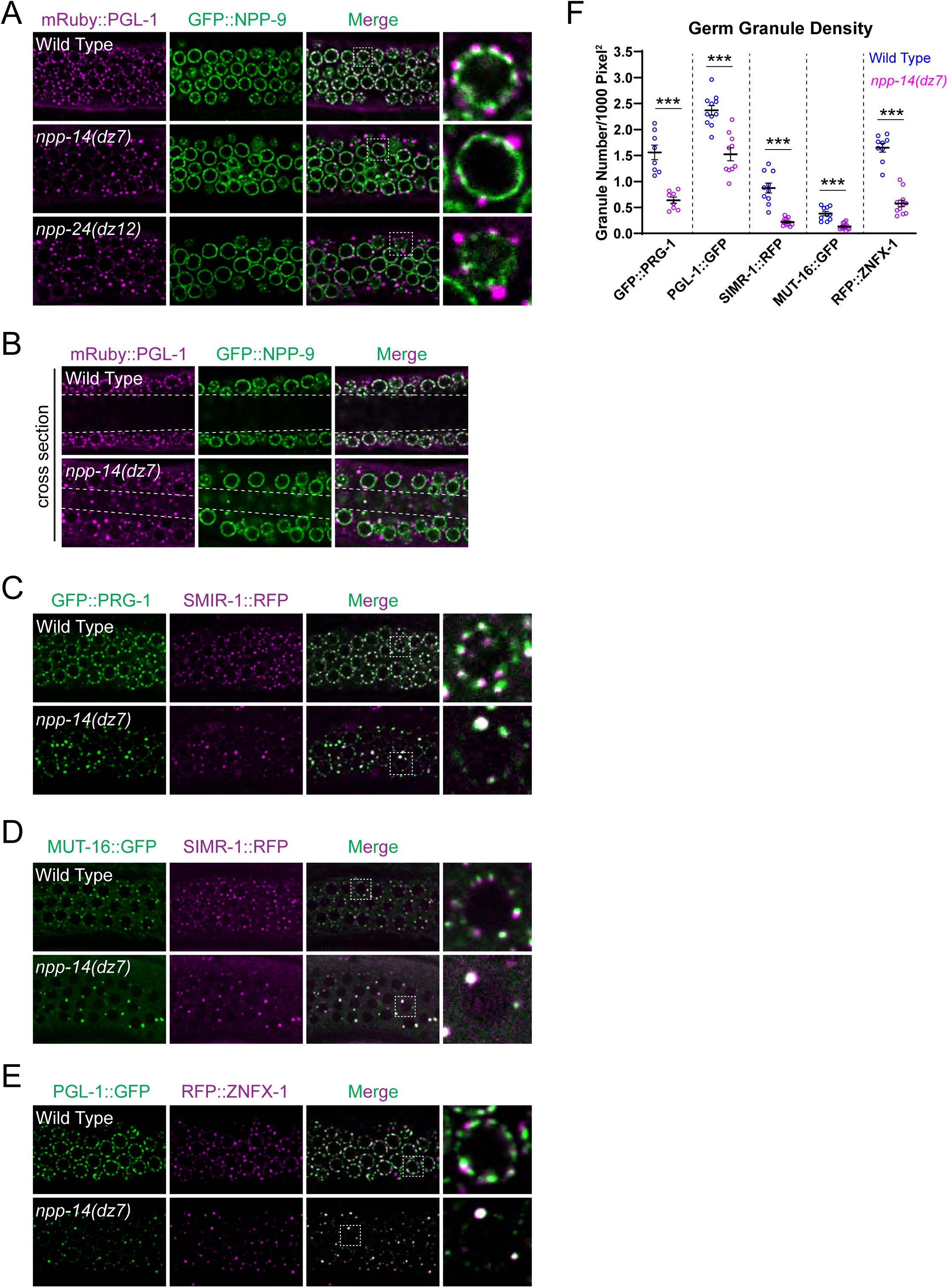
Fewer but enlarged, unorganized germ granules are found in the *npp-14* or *npp-24* mutants. (A-E) Fluorescent micrographs showing the localization of (A-B) mRuby::PLG-1 and GFP::NPP-9, (C) GFP::PRG-1 and SMIR-1::RFP, (D) MUT-16::GFP and SMIR-1::RFP, and (E) PGL-1::GFP and RFP::ZNFX-1, and at the germline nuclei from the pachytene region in the indicated strains. (F) Mean granule density in the indicated strains. Statistical analysis was conducted using a one-tailed Student’s t-test. n=12-15. ***, p<0.001.

While PIWI PRG-1 is enriched in P granules, piRNA downstream factors are enriched in various sub-components of germ granule, including Z granule, SIMR foci, and Mutator foci^10–12^. We therefore examined markers of other sub-compartments of germ granules, including the SIMR foci marker SIMR-1::RFP, the Mutator foci marker MUT-16::GFP, and the Z granule markers RFP::ZNFX-1 or GFP::WAGO-4 in the *npp-14(dz7)* mutant. We found that all examined germ granule markers displayed similar phenotype: fewer but larger perinuclear foci in the *npp-14(dz7)* mutant (Fig. 3C-F, Fig. S3C and S3D). The reduction in the number of foci could not be attributed to decreased protein levels, as quantification of fluorescence intensity or western blot analysis of these germ granule proteins indicated largely unchanged levels (Fig. S3E). Remarkably, unlike in the wild type where germ granules are organized into distinct sub-compartments, the few remaining large germ granules in the *npp-14(dz7)* mutant simultaneously contained proteins from different sub-compartments. This included increased co-localization between P granule markers with SIMR markers, Mutator markers with SIMR-1 markers, P granule markers with Z granule markers, and Z granule with SIMAR-1 markers (Fig. 3C-E and Fig. S3C). Those foci observed in the syncytial cytoplasm contain multiple germ granule proteins and/or the cytoplasmic nucleoporin NPP-9 that we examined (Fig. 3B, Fig. S3F and S3G). Collectively, our observations revealed the critical role of NPP-14 in promoting both the nuclear tethering and organization of germ granules.

### Proximity labeling identifies EPS-1 as a critical germ granule factor that attenuates piRNA silencing

Our observation that NPP-14 and NPP-24 contribute to the organization and nuclear tethering of germ granules prompted us to identify germ granule factors involved in the interactions between NPCs and P granules. Therefore, we conducted proximity labeling and proteomic analyses NPP-14 and PRG-1, components of NPC and P granule, respectively^31^. We compared streptavidin-enriched biotinylated proteins in the strain expressing TurboID::NPP-14 with those in the control strain expressing only cytoplasmic TurboID. Using criteria of 4-fold enrichment and a p-value < 0.05, we identified 184 proteins significantly enriched by TurboID::NPP-14 labeling (Fig. 4A and Supplementary Table 1).

**Figure 4.**
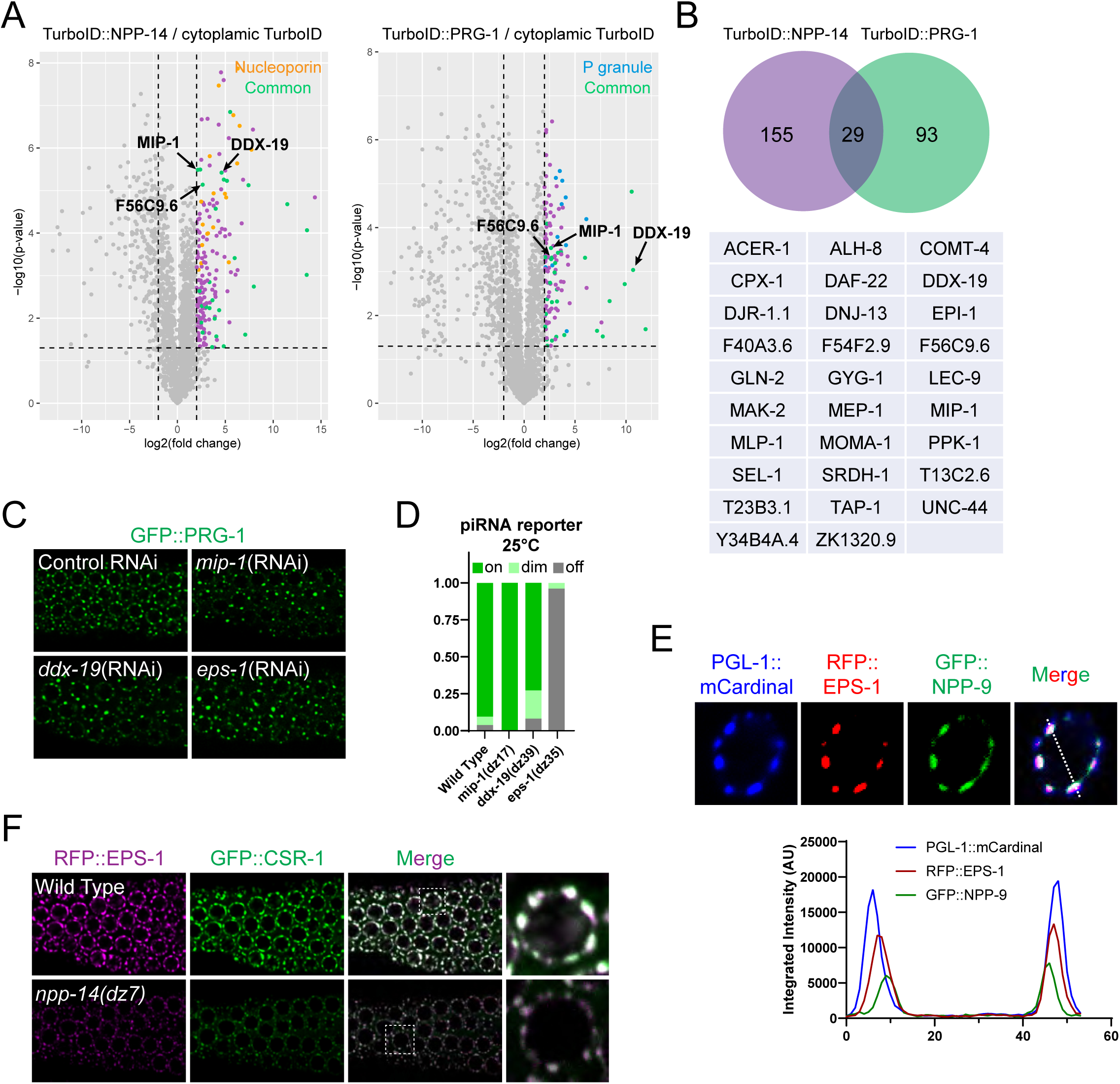
Proximity labeling-mass spectrometry analyses identify EPS-1 as a germ granule factor that negatively regulate piRNA silencing. (A) Volcano plots display the enriched proteins from strains expressing TurboID::NPP-14 and TurboID::PRG-1. Nucleoporins enriched in TurboID::NPP-14 are denoted as orange dots, P granule factors enriched in TurboID::PRG-1 are denoted as blue dots, and proteins enriched in both experiments are depicted as green dots. Statistical significance criteria are log2(Fold change)>2 and p<0.05. (B) A Venn diagram showing the overlap between the enriched proteins from strains expressing TurboID::NPP-14 and TurboID::PRG-1. The overlapped proteins are listed at the bottom. (C) Fluorescent micrographs showing the localization of GFP::PRG-1 in the animals treated with RNAi against the indicated genes. L4440 empty vector was utilized as the RNAi control. (D) Percentage of screened animals showing the GFP expression levels of the piRNA reporter in the indicated strains grown at 25°C. (E) Fluorescent micrographs showing the localization of PGL-1::mCardinal, RFP::EPS-1, and GFP::NPP-9 in the wild type germline nucleus. The line in the merged image indicates the position of the line scan for measuring fluorescent intensity across a single germline nucleus. (F) Fluorescent micrographs showing the localization of RFP::EPS-1 and GFP::CSR-1 in the wild type germline nucleus.

Among these proteins, more than twenty NPC subunits were identified, underscoring the effectiveness of our TurboID-based proteomic analysis (Fig. 4A). Similarly, we identified 122 proteins significantly enriched in the TurboID::PRG-1 expressing strain (Fig. 4A), among which 11 are known P granule proteins. Intriguingly, we found 29 proteins enriched in both TurboID::NPP-14 and TurboID::PRG-1 analyses (Fig. 4B). To determine whether these factors play a role in regulating germ granule formation, we conducted a candidate RNAi screen for some of these genes (Fig. S1B). We discovered that knockdown of *mip-1*, *ddx-19*, or *f59c9.6* resulted in decreased number of perinuclear PRG-1 foci (Fig. 4C). MIP-1/EGGD-1 is a LOTUS domain-containing P granule protein known to promote the assembly and tethering of P granules^31,32^, while DDX-19 is an RNA helicase previously identified as a component of germ granules^18^. We further examined whether *mip-1*, *ddx-19* and *f59C9.6* play a role in piRNA reporter silencing using the null mutants generated by CRISPR-Cas9 editing (Fig. 4D). MIP-1 has been reported to promote piRNA silencing and thus it was not surprising that we observed no enhancement of piRNA reporter silencing in the *mip-1 (dz17)* null mutant animals^33^. *ddx-19 (dz39)* mutant exhibited slightly enhanced silencing of the piRNA reporter but we did not further investigate the role of *ddx-19* (Fig. 4D). Notably, we observed that *f56c9.6* null mutant resulted in enhanced silencing of the piRNA reporter. We designated the *f56c9.6* gene as *eps-1* (<u>e</u>nhanced <u>p</u>iRNA <u>s</u>ilencing-1) due to its mutant *eps-1(dz35)* phenotype.

EPS-1 is an uncharacterized protein that has previously been found to associate with P granule factors^31,34^. To exmamine the localization of EPS-1, we generated a strain expressing PGL-1::mCardinal, RFP::EPS-1, and GFP::NPP-9. Interestingly, RFP::EPS-1 granules localize between P granules and NPCs (Fig. 4E). In addition, we found that EPS-1 co-localize with CSR-1, a D granule enriched factor (Fig. 4F)^36^. D granule was recently reported as a sub-compartment of germ granule that localize between P granule and NPC. Collectively, our imaging analyses of EPS-1 support a model in which EPS-1 is situated between P granules and NPCs to coordinate their interactions.

### EPS-1 acts downstream of NPP-14 but upstream of MIP-1 to regulate germ granule organization

To further investigate the role of *eps-1* in germ granule assembly and organization, we simultaneously monitored two distinct germ granule markers in the *eps-1(dz35)* mutant. Similar to the *npp-14(dz7)* mutant, we observed fewer but larger foci of various granule markers in the *eps-1(dz35)* mutant animals, including PRG-1 foci, ZNFX-1 foci, SIMR-1 foci, and MUT-16 foci (Fig. 5A-C and S4A). Unlike the *npp-14(dz7)* mutant, we did not observe the presence of germ granule factors in the syncytial cytoplasm in the *eps-1(dz35)* mutant (Fig. S4B).

**Figure 5.**
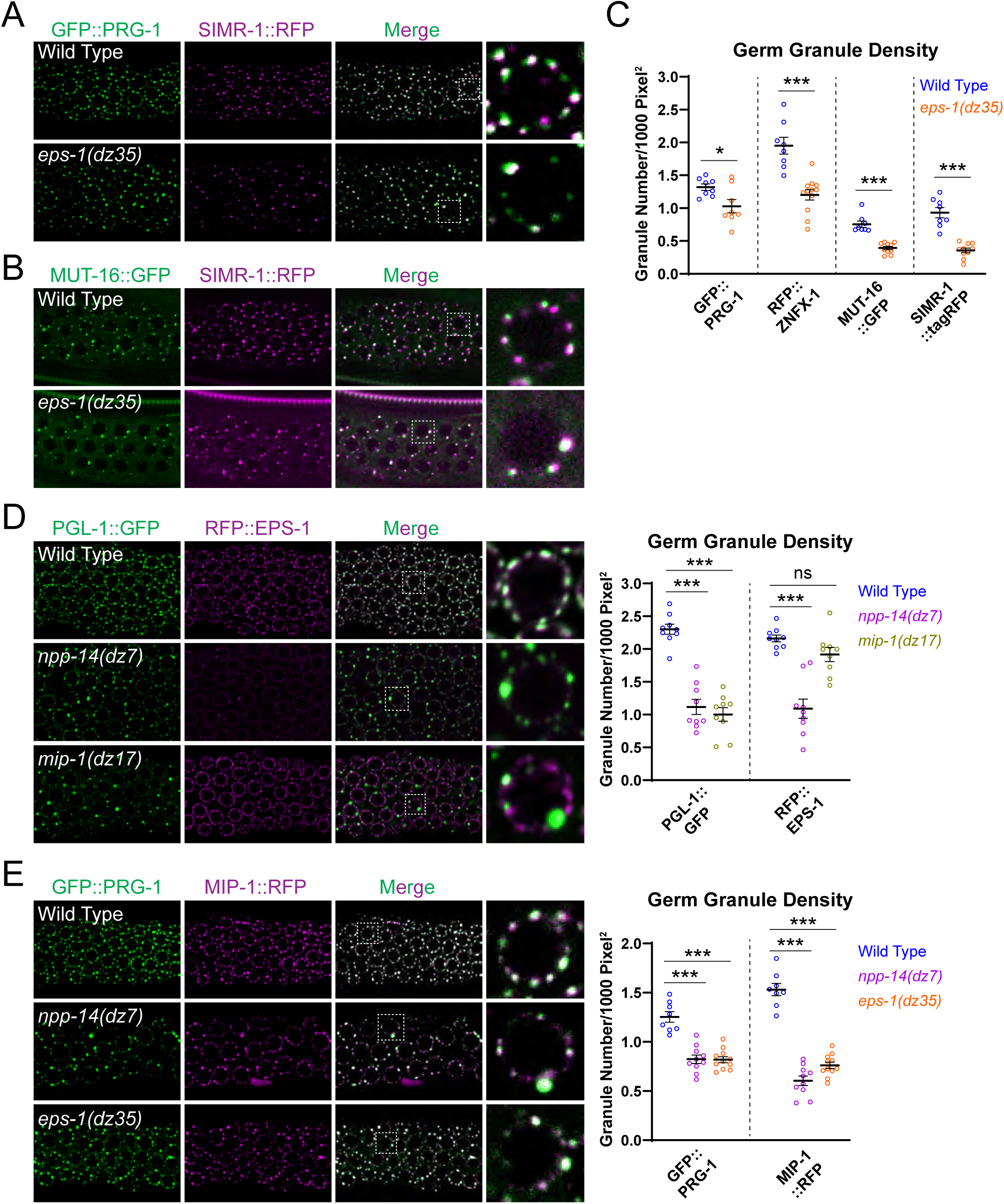
EPS-1 acts downstream of NPP-14 but upstream of MIP-1 in regulating germ granule organization. (A-B) Fluorescent micrographs showing the localization of (A) GFP::PRG-1 and SMIR-1::RFP, (B) MUT-16::GFP and SMIR-1::RFP in the indicated strains. (C) Mean granule density in the indicates strains. Statistical analysis was conducted using a one-tailed Student’s t test. n=12-15. *, p<0.05; ***, p<0.001. (D) Fluorescent micrographs showing the localization of PGL-1::GFP and RFP::EPS-1 in the indicated strains. Comparisons of mean granule density was calculated by one-way ANOVA followed by Dunnett’s correction for multiple comparisons. (E) Fluorescent micrographs showing the localization of GFP::PRG-1 and MIP-1::RFP in the indicated strains. Comparisons of mean granule density was calculated by one-way ANOVA followed by Dunnett’s correction for multiple comparisons.

Since enlarged perinuclear germ granules were found in *mip-1*^31^, *npp-14* and *eps-1* mutants, we next investigated whether EPS-1 is recruited by MIP-1 and NPP-14 to germ granules for regulating germ granule formation. We found that the perinuclear accumulation and protein levels of RFP::EPS-1 were reduced in the *npp-14(dz7)* mutant (Fig. 5D and Fig. S4C). While the localization of PGL-1 was compromised in the *mip-1(dz17)* mutant, neither RFP::EPS-1 enrichment at the perinuclear region nor EPS-1 protein stability were affected in the *mip-1(dz17)* mutant (Fig. 5D and Fig. S4C). These results showed that the abundance and perinuclear localization of EPS-1 were dependent on NPP-14 but not MIP-1. In contrast, the numbers of MIP-1::RFP foci, like other P granule proteins, were reduced in *eps-1(dz35)* or in *npp-14(dz7)* mutants (Fig. 5E), despite the protein levels of MIP-1::RFP not changing in either strain (Fig. S4C). Nonetheless, since there are few residual perinuclear MIP-1 foci and other germ granule marker foci were observed in the *npp-14(dz7)* or in the *eps-1(dz35)* mutants, suggesting that while NPP-14 and EPS-1 contribute to MIP-1 localization, an NPP-14-independent pathway also exists to retain MIP-1 and other germ granule factors at perinuclear regions. Collectively, these results suggest that EPS-1 acts downstream of NPP-14 but upstream of MIP-1 in regulating germ granule organization.

### NPP-14, NPP-24, and EPS-1 prevent the over-production of downstream small RNAs from piRNA targeting sites

piRNAs trigger gene silencing by inducing the production of secondary small RNA, known as WAGO-22G RNA, to promote gene silencing^5^. To investigate how piRNA reporter silencing is enhanced in *npp-14(dz7)*, *npp-24(dz12)*, or *eps-1(dz35)* mutants, we performed small RNA-seq to examine which step of the piRNA-mediated gene silencing pathways was affected. We found that the overall piRNA levels are not changed in *npp-14(dz7)*, *npp-24(dz12)*, and *eps-1(dz35)* mutants (Fig. 6A). In addition, the overall levels of 22G RNAs produced from WAGO-targeted genes were largely unchanged in the *npp-14(dz7)* mutant, although they were slightly reduced in *npp-24(dz12)* and *eps-1(dz35)* mutants (Fig. 6B). Importantly, when we examine specific piRNA targeting sites, such as the piRNA targeting site of the GFP reporter or the endogenous piRNA target *comt-3*^35^, we found that the production of 22G-RNAs was elevated at these sites in *npp-14(dz7)*, *npp-24(dz12)*, and *eps-1(dz35)* mutants (Fig. 6C and 6D). This observation suggests that NPP-14, NPP-24 and EPS-1 restrict the local production of 22G-RNA at these piRNA targeting sites.

**Figure 6.**
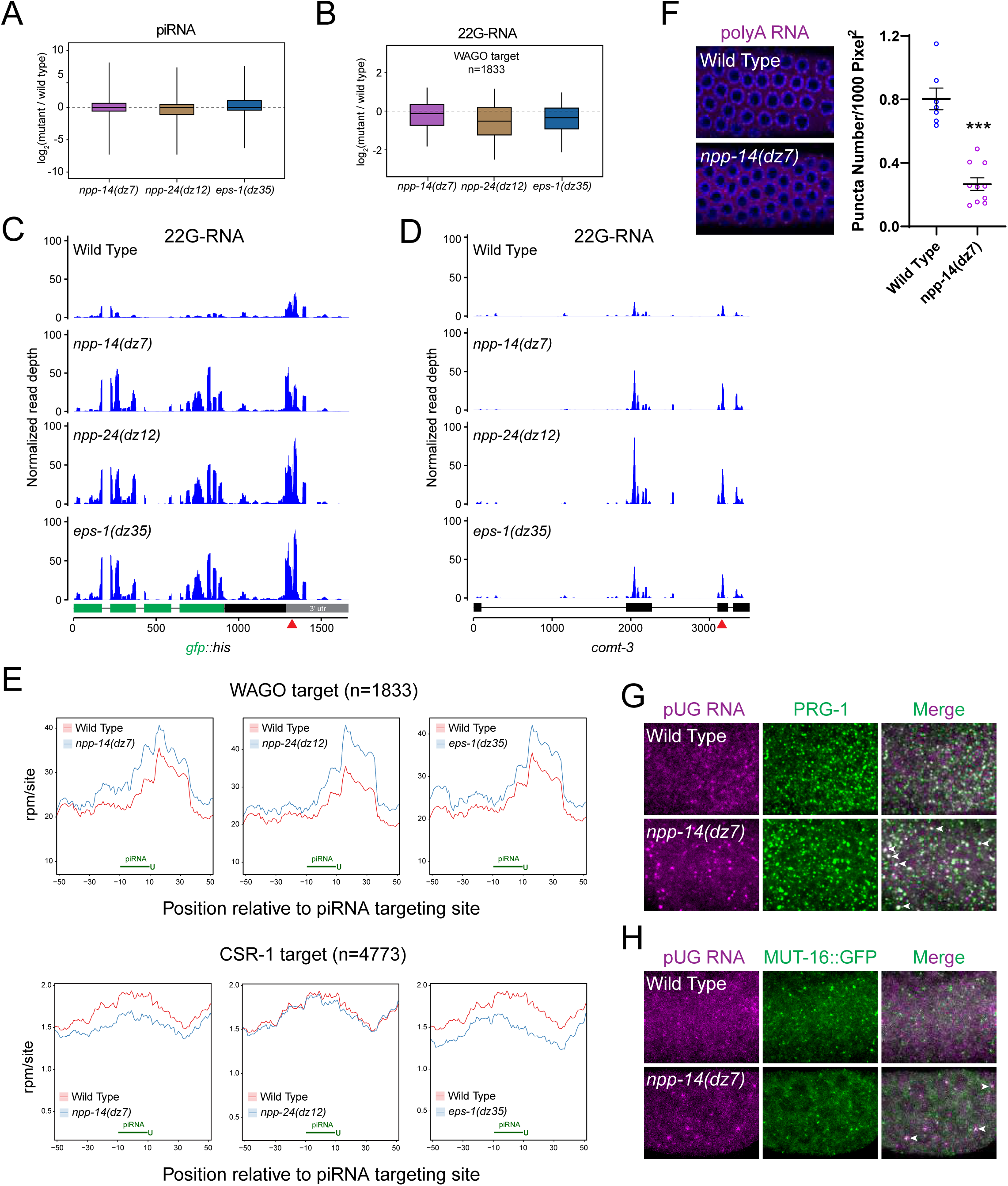
NPP-14, NPP-24, and EPS-1 prevent the over-amplification of secondary small RNAs at piRNA targeting sites (A) Boxplots showing the fold change of piRNA expression level in the indicated strains compared to the wild type. Lines represent median values, boxes represent first and third quartiles, and whiskers display 5th and 95th percentiles. (B) Boxplots showing the fold change of the antisense 22G-RNA, the secondary small RNAs, expression level for WAGO targets in the indicated strains compared to the wild type. Lines represent median values, boxes represent first and third quartiles, and whiskers display 5th and 95th percentiles. (C) Distribution and levels of antisense 22G-RNAs mapped to the piRNA reporter sequences in the indicated strains. The red arrowhead indicates the location of sequences complementary to the endogenous piRNA 21ur-1. (D) Distribution and levels of anti-sense 22G-RNAs mapped to the endogenous piRNA targeting gene *comt-3* sequences in the indicated strains. The red arrowhead indicates the location of predicted piRNA targeting site. (E) Density of 22G-RNAs within a 100-nt window around predicted piRNA targeting sites in the indicated strains. Computed by summing 22G-RNA density per piRNA targeting site in WAGO targets (top) or CSR-1 targets (bottom). The plots are centered on the 10th nucleotide of piRNAs. (F) Fluorescent micrographs of pachytene nuclei in fixed adult gonads hybridized with a fluorescent polyT probe complementary to polyA RNA. Nuclear DNA was stained with DAPI (Blue). (G-H) Fluorescent micrographs of pachytene nuclei of fixed adult gonads hybridized with a fluorescent probe complementary to pUG RNA and immunofluorescence stained with (G) PRG-1 or (H) GFP antibodies in the indicated strains. The arrowheads indicate colocalized pUG RNA foci with PRG-1 foci or with MUT-16::GFP foci.

We next examined the production of 22G-RNAs at piRNA binding sites on a transcriptome-wide scale^37^. Specifically, we compared the levels of 22G-RNAs from WAGO target genes (germline-silenced genes) or from CSR-1 target genes (germline-expressed genes) around predicted piRNA targeting sites^38–40^. We found that 22G-RNA levels were increased at piRNA binding sites of WAGO target genes in *npp-14(dz7), npp-24(dz12)*, and *eps-1(dz35)* mutants (Fig. 6E). In contrast, 22G-RNA levels around the piRNA binding sites of CSR-1 target genes were decreased or unchanged in the *npp-14(dz7)*, *npp-24(dz12)*, and *eps-1(dz35)* mutants (Fig. 6E).

We next investigated whether these elevated 22G-RNAs at WAGO target genes were indeed loaded onto WAGO Argonautes, such as WAGO-1 and HRDE-1. We performed WAGO-1, HRDE-1, and CSR-1 IP and small RNA-seq to measure the levels of small RNA associated with these Argonaute proteins in wild type and in the *npp-14(dz7)* mutant. We found that WAGO-1 and HRDE-1 bound 22G-RNAs, but not CSR-1 bound 22G-RNAs, were increased at the piRNA reporter mRNA (Fig. S5A). Notably, WAGO-1 22G-RNAs were increased across the piRNA reporter mRNA, whereas HRDE-1 22G-RNAs were increased more locally at the piRNA targeting site in the *npp-14*(dz7) mutant).

Transcriptome-wide analyses showed increased levels of WAGO-1 and HRDE-1 associated 22G-RNAs at piRNA binding sites of WAGO targeting genes and decreased levels of CSR-1 associated 22G-RNAs at piRNA targeting sites of CSR-1 targets (Fig. S5B). Taken together, our results suggest that NPP-14, NPP-24, and EPS-1 prevent the overproduction of WAGO 22G-RNAs from the piRNA binding sites.

### Enlarged germ granules in the *npp-14* mutant are enriched with prominent pUG RNA foci

As describe earlier, P granule, a sub-compartment of germ granule, are known to be the major sites of mRNA export in the germline of *C. elegans*^18^. Since fewer and larger germ granules are found in the *npp-14(dz7)* mutant, we investigated whether mRNA export and localization is affected. We performed smFISH against PolyA RNAs in the *npp-14(dz7)* mutant. In wild-type animals, in addition to cytoplasmic signal, PolyA RNAs were accumulated at perinuclear foci (Fig. 6F). In the *npp-14(dz7)* mutant, while PolyA RNAs were also present in the cytoplasm, the number of perinuclear foci was significantly reduced. This result indicates that mRNAs are either accumulated or exported at fewer perinuclear foci in the *npp-14(dz7)* mutant (Fig. 6F).

Once piRNA recognizes its mRNA targets, additional piRNA pathway factors RDE-8 and RDE-3/MUT-2 are recruited to cleave and to add polyUG RNA tails (pUG RNAs) on the mRNA targets, respectively^18,43^. pUG RNAs act as a key signal that recruits RNA dependent RNA polymerase to produce downstream WAGO 22G-RNAs^18^. As WAGO 22G-RNAs are over-produced from piRNA targeting sites in the *npp-14(dz7)* mutant, we asked whether pUG RNAs are accumulated in perinuclear germ granules in the *npp-14(dz7)* mutant. By using smFISH and immunostaining, we found that more prominent pUG RNA foci were found in the *npp-14(dz7)* mutant than in wild-type animals, and many of these large pUG RNA foci were co-localized with PIWI PRG-1 or Mutator foci markers MUT-16 (Fig. 6G and 6H). These results show those enlarged germ granule found in *npp-14* mutant animals are enriched with pUG RNAs. Collectively, our imaging analyses and small RNA sequencing data where the piRNA and its mRNA targets were both enriched in those large perinuclear germ granules found in the *npp-14(dz7)* mutant, leading to the formation of larger pUG RNA foci and an increased production of secondary WAGO 22-RNAs from piRNA targeting sites.

### Impairment of RNAi silencing and inheritance in the *npp-14* mutant

piRNAs and siRNAs share common downstream WAGO 22G-RNA pathway factors to trigger gene silencing, and previous studies have suggested that these two pathways can compete for downstream gene silencing machinery^6^. In the *prg-1* mutant animals, which lose all piRNAs, RNAi inheritance is greatly enhanced^6^. Given the enhanced piRNA-mediated gene silencing found in the *npp-14(dz7)* mutant, we wondered whether the efficiency of RNAi may be affected. To compare the germline RNAi efficiency, we assessed the ratio of dead embryos upon *pos-1* RNAi. *Pos-1* is a maternally expressed gene required for embryonic viability^41^. While the treatment of undiluted *pos-1* dsRNA-expressing bacteria led to nearly 100% lethality in both wild-type and the *npp-14(ok1389)* null mutant animals, treatment with diluted *pos-1* dsRNA-expressing bacteria led to a smaller portion of dead embryos in the *npp-14(ok1389)* null mutant than in wild-type animals. (Fig. S6A). This result suggests that the RNAi efficiency is compromised in the *npp-14(ok1389)* mutant.

Proper germ granule organization has been linked to RNAi inheritance^10,42,44^. We next asked whether is compromised in the *npp-14* mutant. In these experiments, we treated a GFP::histone-expressing strain with GFP dsRNA-expressing bacteria for one generation^44^. The animals were then treated with bleach to obtain their embryos and to avoid bacterial contamination across generations. The embryos of each following generation were cultured on control food to determine the ratio of GFP-silenced animals (Fig. 7A). We found that GFP RNAi treatment could completely silence GFP::histone signals in both the wild type and the *npp-14(dz7)* mutant animals. Notably, in the each of the following three generations on control food, while 100%, 73%, and 55% of wild-type animals still exhibited no GFP signals, only 80%, 20%, and 16% of the *npp-14(dz7)* mutant animals exhibited no GFP signals (Fig. 7B and 7C). Additionally, we sequenced small RNAs and compared the abundance of 22G-RNAs targeting GFP mRNAs. We found that those 22G-RNAs targeting GFP mRNAs were slightly decreased in the *npp-14(dz7)* mutant compared to wild type in the P0 generation but were significantly more decreased in the first generation after removal of the RNAi bacteria (Fig. 7D). Collectively, our data show that the efficiency of RNAi and RNAi inheritance were compromised in the *npp-14* mutant animals. Our results highlight the importance of NPP-14 in maintaining a balanced gene silencing program between siRNA and piRNA in germ cells (Fig. 7E).

**Figure 7.**
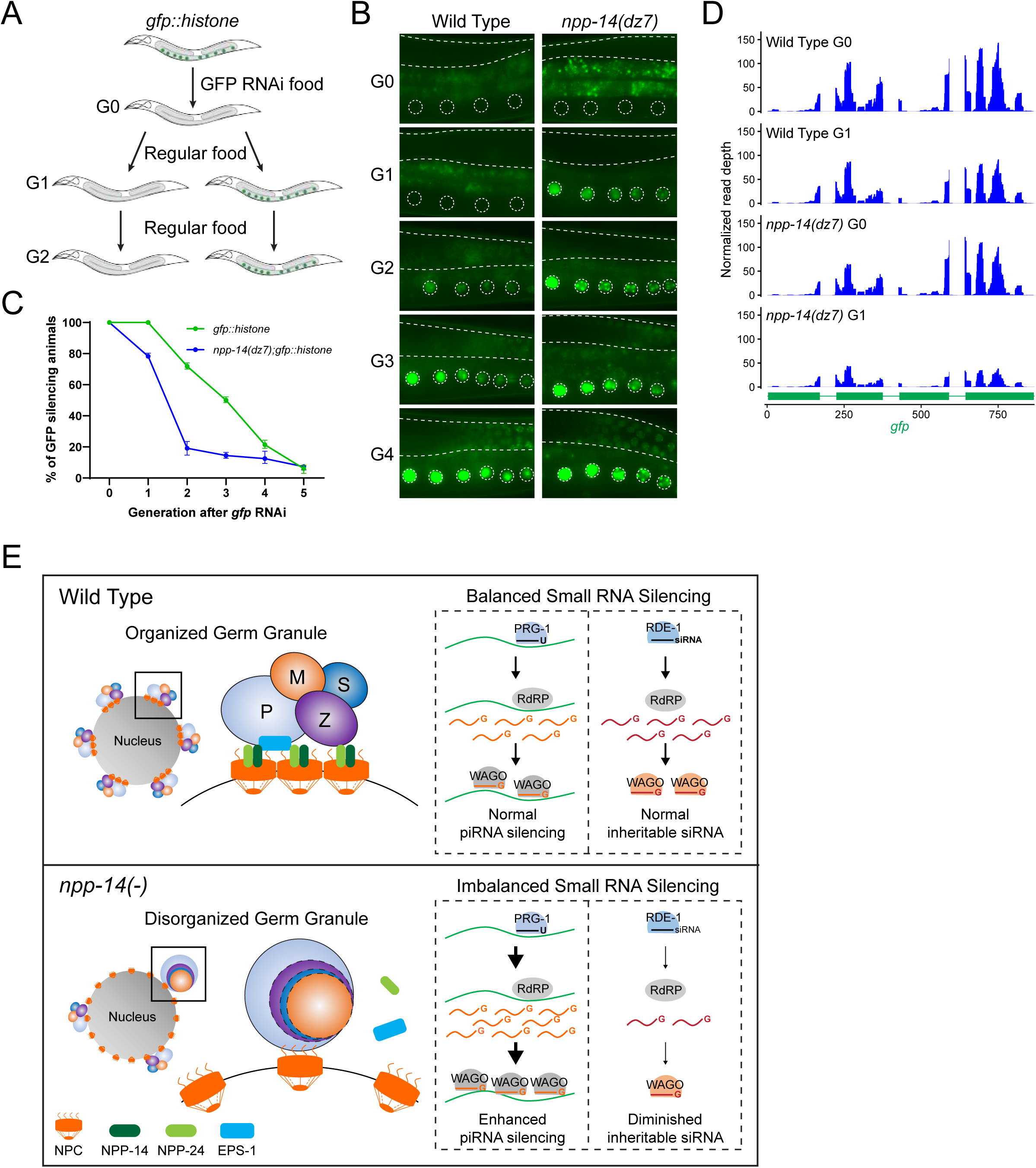
RNAi inheritance is compromised in the *npp-14* mutant. (A) A schematic illustrating the RNAi assay with a GFP::histone reporter to assess the mutant’s ability to initiate RNAi and to inherit gene silencing over generations. (B) Representative fluorescent micrographs showing the GFP::histone reporter in the RNAi-treated generation (G0) or the subsequent generations (G1 to G4) in the indicated strains. Dashed lines and circles denote the germline and maturing oocyte nuclei, respectively. Note that the bright signals observed outside of dashed areas are autofluorescence signals originating from gut granules. (C) Percentage of screened animals showing GFP reporter silencing in the indicated strains. Data points represent the mean, with error bars indicating the standard error of the mean. Distributions represent data collected from three independent experiments. (D) Distribution and levels of the anti-sense 22G-RNAs mapped to GFP coding sequences in the indicated strains from the RNAi-treated generation G0 and the subsequent generation G1. (E) A model depicting the role of NPP-14, NPP-24 and EPS-1 in germ granule organization and its role in balancing small RNA gene silencing pathways.

## DISCUSSION

In this study, we examined factors regulating gene silencing and germ granule-NPC interactions, uncovering the pivotal roles of two nucleoporins, NPP-14 and NPP-24, in germ granule organization and in regulation of piRNA silencing. The enhanced silencing of piRNAs observed in the *npp-14(dz7)* mutant was accompanied by fewer but enlarged perinuclear germ granules, fewer perinuclear mRNA foci, enlarged pUG RNA foci, and the overproduction of secondary WAGO 22G-RNAs at piRNA targeting sites. While the exact mechanism underlying the enhancement of piRNA silencing remains unknown, these observations support a model of how aberrant germ granule enhance piRNA silencing; since the mRNAs are accumulated at fewer perinuclear foci in the *npp-14(dz7)* mutant, where PRG-1 and downstream factors are concentrated, the enrichments of mRNAs and PRG-1 in fewer foci can lead to higher local concentrations that lead to enhance piRNA binding to its mRNA target. As a result, more pUG RNAs are being produced, leading to over-amplification of secondary 22G-RNAs at piRNA targeting sites and enhanced piRNA silencing.

Our study also reveals the key role of NPP-14 and NPP-24 EPS-1 in NPC-germ granule interactions to ensure proper germ granule organization and to restrict piRNA silencing. The exact mechanism of how NPP-14, NPP-24, and EPS-1 contribute to the NPC-germ granule interaction will be an area for further investigation in the future. Nonetheless, we showed that EPS-1 is positioned between P granules and NPCs and it acts downstream of NPP-14 but upstream of MIP-1 to regulate germ granule assembly. Notably, EPS-1 carries multiple intrinsic disordered regions and NPP-14 carries phenylalanine-glycine (FG) repeat. As intrinsic disordered regions and FG repeats are present in multiple germ granule factors and are known to be critical for germ granule assembly, they could play a critical role in establishing germ granule-NPC interaction. The weakened germ granule-NPC interaction may therefore lead to the dissociation of germ granule to into syncytial cytoplasm and fewer perinuclear germ granules.

The enhanced piRNA silencing but compromised siRNA silencing observed in *npp-14* mutant suggest a critical function of germ granule-NPC interactions in balancing siRNA and piRNA gene silencing, two major small RNA-mediated genome defense pathways in animals^3^. Notably, only the piRNA Argonaute PRG-1, but not the siRNA Argonaute RDE-1, is known to be enriched in germ granules^45,46^. As different foreign/selfish RNAs may be present at distinct location in the cells, such as RNAs transcribed from transposable elements in the nucleus or the virual dsRNA in the cytoplasm, our study raise an interesting model where the compartmentalization of small RNA gene silencing factors may have evolved to optimize the genome defebse mechanisms against various foreign/selfish RNAs.

## METHODS

### Caenorhabditis elegans strains

Animals were at either 20 °C or 25 °C cultured on standard nematode growth media (NGM) plates seeded with the *E. coli* OP50 strain. The Bristol strain N2 was utilized as the standard wild-type strain. Details of the strains employed in this study can be found in Supplementary Table 2.

### Brood size

L4 stage hermaphrodites (P0) were individually placed onto freshly seeded NGM plates and allowed to grow for 24 hours at either 20 °C or 25 °C. P0 adults were transferred to new NGM plates every 24 hours until they no longer laid eggs. All the F1 progeny were counted. The brood size of each P0 animal is the total sum of F1 progeny across all plates where the P0 animal laid eggs. At least 8 individual P0 animals were used to score the brood size for each strain.

### RNAi and RNAi inheritance assay

*E. coli* HT115 (DE3) strains expressing dsRNAs against target genes were obtained from *C. elegans* RNAi collection derived from the *C. elegans* ORFeome Library v1.1 (Horizon Discovery) or homemade strains. To prepare the RNAi bacterial cultures, Luria broth supplemented with 100 μg/mL ampicillin was inoculated with the respective bacterial strain and incubated overnight at 37°C. The cultures were then seeded onto NGM plates containing 100 μg/mL ampicillin and 1 mM IPTG and incubated at room temperature for 1 to 2 days to induce dsRNA expression. RNAi experiments were conducted at either 20°C or 25°C by transferring synchronized L1s or L4s onto NGM plates seeded with RNAi bacteria. Adults (transferred at the L1 stage) or progeny adults (transferred at the L4 stage) were subjected to imaging and phenotype scoring. For the pos-1 RNAi treatment, pos-1 RNAi bacteria were diluted with control RNAi bacteria expressing empty vector L4440.

For the RNAi inheritance assay, synchronized L1 animals of indicated genotypes were exposed to bacteria expressing gfp dsRNA. F1 embryos were collected by hypochlorite/NaOH bleach treatment and subsequently cultured on NGM plates seeded with *E. coli* OP50. GFP expression in both the parental generation and progeny were imaged and scored. The silenced GFP reporter was characterized by the absence of GFP or dim GFP expression, while the desilenced GFP reporter was characterized by robust GFP expression. Images were captured using a Zeiss Axio Imager M2 compound microscope with a Plan-Apochromat 40x/1.4 Oil objective.

### CRISPR/Cas9-mediated gene editing and transgenes

CRISPR/Cas9-mediated gene editing experiments were conducted using Cas9/sgRNA ribonucleoprotein (RNP) strategies. The sgRNAs for CRISPR/Cas9 gene editing were transcribed *in vitro* using the HiScribe T7 Quick High Yield RNA Synthesis Kit (New England Biolabs, E2050S) and purified with the Monarch RNA Cleanup Kit (New England Biolabs, T2040L). The 3xFLAG/GFP/tagRFP tag sequence (short donor) and a tag sequence containing short 5′ and 3′ homology sequences (∼40 nt) (long donor) were amplified by PCR. The purified 1.2 µg of short donor and 1.2 µg of long donor products were subjected to a denaturation and renaturation process (95 °C for 2 min, 85 °C for 10 s, 75 °C for 10 s, 65 °C for 10 s, 55 °C for 1 min, 45 °C for 30 s, 35 °C for 10 s, 25 °C for 10 s, and held at 4 °C) to generate hybrid DNA donor templates. The final concentrations of injection mixture were as follows: 250 ng/µL of Alt-R S.p. Cas9 Nuclease V3 (Integrated DNA Technologies), 200 ng/µL of *in vitro* synthesized sgRNA, 200 ng/µL of hybrid DNA donor and 40 ng/µL of co-injection marker pRF4. Cas9 and sgRNA were mixed and incubated at 37 °C for 10 min before adding other components. The injection mixtures were microinjected into the gonads of adult *C. elegans*. The F1 roller progeny were individually collected and genotyped by PCR to identify the desired transgenes.

Mos1-mediated single copy insertion (MosSCI) was used to generate the *pie-1p::turboID::3xha* transgenic strain. Final concentrations of injection mixture were as follows: 50 ng/µL of pCFJ601 (eft-3p::transposase), 2.5 ng/µL of pCFJ90 (myo-2p::mCherry), 5 ng/µL of pCFJ104 (myo-3p::mCherry) and 50 ng/µL of pDZ615 (pie-1p::turboID::3xha::pie-1 3’utr in pCFJ150). EG8081 (unc-119(ed3) III; oxTi177 IV) was used for injection. Two injected animals were placed on each seeded NGM plate and cultured at 25°C. Plates were screened for animals exhibiting normal movement but lacking the red co-injection markers.

### Fluorescence microscopy and image processing

GFP- and RFP-tagged fluorescent proteins were observed in living nematodes by mounting young adult animals on 2% agarose pads with M9 buffer (22 mM KH_2_PO_4_, 42 mM Na_2_HPO_4_, 86 mM NaCl) supplemented with 50 mM levamisole. Images were captured using either a Zeiss LSM800 confocal microscope with a Plan-Apochromat 63x/1.4 Oil DIC M27 objective or a Zeiss Axio Imager M2 compound microscope with a Plan-Apochromat 40x/1.4 Oil objective. Single slice images of the gonads were used to define Regions of Interest (ROIs), and areas of the ROIs were measured. Fluorescence thresholds were set manually to identify fluorescent puncta within the ROIs. Granule density was calculated by dividing the number of fluorescent puncta within an ROI by the area of the ROI. Integrated fluorescence intensity within the ROIs were measured. Quantitative analysis was based on data from 10 to 15 gonads per strain. Statistical analysis of the data was performed using student’s t-test or one-way ANOVA followed by Dunnett’s correction for multiple comparisons.

### Western blotting

Lysates were prepared from 100 synchronized young adult worms. The worms were boiled in boiling buffer (100 mM Tris-HCl (pH 6.8), 8% SDS, 20 mM β-mercaptoethanol) for 10 minutes and then in 1x SDS loading buffer for an additional 5 minutes. Proteins were separated by standard SDS-PAGE and transferred to PVDF membranes (Roche) using the Trans-Blot Turbo Transfer System (Bio-Rad). The membranes were blocked in 5% skimmed milk in TBST (20 mM Tris-HCl pH 7.4, 150 mM NaCl, and 0.05% Tween 20) for 1.5 hours at room temperature and then incubated with primary antibodies overnight at 4 °C. After five washes with TBST buffer, membranes were incubated with secondary antibodies for 1.5 hours at room temperature then washed five times with TBST. ECL substrates were used for detection of protein bands and imaged were obtained by a Tanon 5200 Chemiluminescent Imaging System (Tanon). The antibodies used in this study were anti-GFP (Santa Cruz, sc-9996) at 1:500 dilution, anti-HA (Cell Signaling Technology, 3724S) at 1:1000 dilution, anti-FLAG (Sigma-Aldrich, F1804) at 1:1000 dilution, anti-α-Tubulin (Sigma-Aldrich, A11126) at 1:1500 dilution, HRP-conjugated Goat Anti-Mouse IgG (H+L) (Proteintech, SA00001-1) at 1:5000 dilution, and HRP-conjugated Goat Anti-Rabbit IgG (H+L) (ZSGB-Bio, ZB-2301) at 1:5000 dilution.

### TurboID-based proximity labeling

Approximately 15,000 synchronized young adults were washed three times with M9 buffer and subsequently incubated in M9 buffer supplemented with 1 mM biotin for 2 hours at 20 °C. After incubation, the worms underwent three washes in M9 buffer followed by one wash in pre-cooled ddH_2_O. Animals were flash-frozen in liquid nitrogen and stored at -80 °C until further use.

Two volumes of RIPA buffer (50 mM Tris-HCl pH 7.5, 150 mM NaCl, 0.5% sodium deoxycholate, 1% NP-40) supplemented with 1x protease inhibitor cocktail without EDTA (Promega, G6521) were added to one volume of packed worms, which were then ground in a glass grinder 8-10 times on ice within a 10-minute timeframe. Lysates were clarified by centrifugation at 15,000 rpm, 4 °C, for 15 minutes. The resulting supernatants were desalted using Zeba spin desalting columns 7K MWCO (Thermo Scientific, 89890), followed by incubation with Dynabeads MyOne streptavidin C1 (Invitrogen, 65002) at room temperature for 30 minutes on a rocker. Beads were washed five times with RIPA buffer and twice with 1x PBS buffer. After removing excess buffer, beads were stored at -80°C for mass spectrometry analysis.

### Mass spectrometry analysis

Streptavidin beads were boiled in UA buffer (8M urea, 150 mM Tris-HCl, pH 8.0) supplemented with 100 mM dithiothreitol for 5 minutes. The beads were then washed three times with 50 mM ammonium bicarbonate, then incubated with 50 mM iodoacetamide in the dark at room temperature for 30 minutes. After incubation, beads were mixed with 6 µg of trypsin (Promega, V5113) in 40 µL of 50 mM ammonium bicarbonate and digested at 37 °C for 16-18 hours. The supernatant containing the released peptides was collected. The peptides were desalted using C18 StageTip for subsequent LC-MS/MS analysis. LC-MS/MS analysis was conducted using a Q Exactive Plus mass spectrometer coupled with an Easy 1200 nLC system (Thermo Fisher Scientific). Initially, peptides were loaded onto a trap column (100 μm x 20 mm, 5 μm, C18, Dr. Maisch GmbH) with buffer A (0.1% formic acid in water). For reverse-phase high-performance liquid chromatography (RP-HPLC) separation, an EASY-nLC system (Thermo Fisher Scientific, Bremen, Germany) equipped with a self-packed analytical column (75 μm x 150 mm; 3 μm ReproSil-Pur C18 beads, 120 Å, Dr. Maisch GmbH) was used at a flow rate of 300 nL/min. Mobile phase A consisted of 0.1% formic acid in water, while mobile phase B was 0.1% formic acid in 95% acetonitrile. Peptides were eluted over 120 minutes using a linear gradient of buffer B. Mass spectrometry data was acquired employing a data-dependent top20 method that dynamically selected the most abundant precursor ions from the survey scan (300-1800 m/z) for higher-energy collisional dissociation (HCD) fragmentation. Peptide recognition mode was enabled during the analysis, with a lock mass of 445.120025 Da utilized for internal mass calibration. Full MS scans were acquired at a resolution of 70,000 at m/z 200, and MS/MS scans at a resolution of 17,500 at m/z 200. The maximum injection time was set to 50 ms for both MS and MS/MS scans. The normalized collision energy was 27, and the isolation window for precursor ion selection was set to 1.6 Th. Dynamic exclusion was applied with a duration of 60 seconds. The mass spectrometry database search software MaxQuant 1.6.1.16 was used with the following protein database from Uniprot Protein Database: species C. elegans uniprot-C. elegans [6239]-27419-20210222.fasta. The database search results were filtered and exported with <1% false discovery rate (FDR) at peptide-spectrum-matched level and protein level, respectively.

### RNA isolation and small RNA sequencing

Total RNA was extracted from immunoprecipitated samples or whole animals of approximately 100,000 synchronized young adults using TRIzol reagent (Invitrogen, AM9738) following a standard protocol. Small (<200 nt) RNAs were then enriched using the mirVana miRNA Isolation Kit (Ambion, AM1561). In brief, 80 μL of total RNA (<1,000 μg) was mixed with 400 μL of mirVana lysis/binding buffer and 48 μL of mirVana homogenate buffer, followed by a 5-minute incubation at room temperature. Subsequently, 176 μL of 100% ethanol was added, and the mixture was centrifuged at 2500×g for 4 minutes at room temperature to pellet large (>200 nt) RNAs. The supernatant containing small (<200 nt) RNAs was transferred to a new tube, and these small RNAs were precipitated with pre-cooled isopropanol at -70°C. After centrifugation at 20,000×g at 4°C for 30 minutes, the RNA pellet was washed once with 70% pre-cooled ethanol and dissolved in nuclease-free water. 10 μg of small RNAs were fractionated on a 16% PAGE gel containing 7 M urea, and RNA within the range of 18 nt to 40 nt was excised from the gel. RNA was extracted by soaking the gel in 700 µL of NaCl TE buffer (0.3 M NaCl, 10 mM Tris-HCl, 1 mM EDTA, pH 7.5) overnight. The supernatant was collected through a gel filtration column, and RNA was precipitated with isopropanol, washed with 70% ethanol, and resuspended in 15 μL of nuclease-free water. The RNA samples underwent treatment with RppH (New England Biolabs, M0356S) to convert 22G-RNA 5′ triphosphates to monophosphates. RNA was then concentrated using the RNA Clean and Concentrator-5 Kit (Zymo Research, R1015).

Small RNA libraries were prepared using the NEBNext Multiplex Small RNA Sample Prep Set for Illumina-Library Preparation (New England Biolabs, E7300) according to the manufacturer’s protocol. NEBNext Multiplex Oligos for Illumina Index Primers (New England Biolabs, E7335) were used for library preparation. The libraries were sequenced using the Illumina NovaSeq 6000 system.

### RNA immunoprecipitation sequencing (RIP-seq)

Approximately 100,000 synchronized young adults were used for RIP-seq. Worms were washed three times with M9 buffer. Worm pellets were resuspended in an equal volume of IP buffer (20 mM Tris-HCl pH 7.5, 150 mM NaCl, 2.5 mM MgCl_2_, 1 mM dithiothreitol, 0.5% NP-40, 80 U/mL RiboLock RNase Inhibitor (Thermo Scientific, EO0381), 1x protease inhibitor cocktail without EDTA (Promega, G6521)). The pellets were ground in a glass grinder 8-10 times within a 10-minute timeframe. Lysates were clarified by centrifugation at 15,000 rpm, 4 °C, for 15 minutes. Supernatants were incubated with Anti-Flag Magnetic Beads (MedChemExpress, HY-K0207) at 4 °C for 1.5 hours on a rocker. Beads were washed five times with IP wash buffer (20 mM Tris-HCl pH 7.5, 150 mM NaCl, 2.5 mM MgCl_2_, 1 mM dithiothreitol, 0.5% NP-40), followed by two washes with 1x PBS buffer. Total RNA was extracted using TRIzol reagent (Invitrogen, AM9738) following standard protocols. Small RNA libraries for RNA-seq were prepared as described above and sequenced using an Illumina NovaSeq 6000 system.

### Small RNA sequencing data analysis

Fastq reads were trimmed using custom Perl scripts . Subsequently, the trimmed reads were aligned to the *C. elegans* genome build WS230 using Bowtie version 1.3.1^47^ with options -v 0 -best -strata. Following alignment, reads ranging from 17 to 40 nucleotides in length were intersected with various genomic features (rRNAs, tRNAs, snoRNAs, miRNAs, piRNAs, protein-coding genes, pseudogenes, and transposons) using BEDTools intersect. Sense and antisense read mappings to specific genomic features were quantified and normalized to reads per million (RPM) by scaling read counts based on total mapped reads, excluding those aligning to structural RNAs (rRNAs, tRNAs, snoRNAs). To account for reads mapping to multiple loci, the read count was adjusted by dividing it by the number of loci where each read perfectly aligned. Specifically, only reads matching the sense annotation without overlap were considered for miRNAs and piRNAs. Reads classified as 22G-RNAs were defined as 21 to 23 nucleotide-long reads with a 5′ G that aligned to protein-coding genes, pseudogenes, or transposons. The resulting RPM values were utilized in downstream analyses, conducted using custom R scripts in R version 4.3.3, leveraging packages including ggplot^48^, reshape^49^, ggpubr^50^, and dplyr^51^.

Metagene profiles across gene lengths were generated by analyzing the depth at each genomic position using 21 to 23 nucleotide-long small RNA reads starting with a 5′ G using BEDTools genomecov^52^. Metagene profiles relative to piRNA targeting sites were computed as the mean normalized reads per million at each nucleotide position using the indicated 22G-RNA reads. piRNA targeting sites were identified based on stringent rules for piRNA targeting within the indicated group of transcripts according to previously published guidelines for piRNA target prediction. Groups of genes were then plotted based on the cumulative normalized depth across a 100-nt window around predicted piRNA target sites. All custom scripts can be found at the supplemental data file 1.

### Single molecule fluorescence in situ hybridization (smFISH) and PRG-1 and MUT-16::GFP immunohistochemistry

PolyA or polyUG RNA smFISH was performed on dissected adult gonads. Young adult worms were washed with M9 buffer and transferred onto a glass slide containing Dissection Buffer (1x PBS, 0.1% Tween-20, 0.25 mM levamisole, 1 mM EDTA). Worms were dissected using a 1 mL syringe needle, and then transferred to individual 1.5 mL microcentrifuge tubes. Worms were fixed by incubating with 1 mL of Fix Solution (1x PBS, 3.7% formaldehyde, 0.1% Tween-20) for 30 minutes at room temperature on a rocker. After fixation, worms were washed once with 1 mL of PBST (1x PBS, 0.1% Tween-20) and then incubated for 10 minutes with 1 mL of 1x PBS containing 0.1% Triton X-100 at room temperature on a rocker to permeabilize the cells. Following permeabilization, worms were washed twice with 1 mL of PBST, resuspended in 1 mL of 70% ethanol and incubated at 4 °C for 16-18 hours for further processing.

For smFISH combined with immunohistochemistry, dissected worms were washed once with Antibody Wash Buffer (1x PBS, 1 mM EDTA, 0.1% Tween-20) and 50 μL of anti-PRG-1 (custom-made anti-rabbit antibody provided by Dr. Craig Mello lab) or anti-GFP (Santa Cruz, sc-9996) suspended at 1:200 dilution in Antibody Suspension Buffer (1 x PBS, 1 mM EDTA, 0.1% Tween-20, 20 mg/mL BSA, 2 mM vanadyl ribonucleoside complex) was added. Samples were shaken at 850 rpm overnight at 4 °C, washed twice with Antibody Wash Buffer, and 50 μL of anti-Rabbit Alexa488 (Jackson Labs, 711-547-003) suspended at 1:400 dilution in Antibody Suspension Buffer was added. Samples were shaken at 850 rpm for 2 h at room temperature and washed once with Antibody Wash Buffer and once with 2× SSC.

1.32 µL of the 2.5 µM polyA or polyUG RNA smFISH probe (Sangon Biotech) was added into fresh HB buffer (297 µL Stellaris RNA FISH hybridization buffer (LGC Biosearch Technologies, SMF-HB1-10) and 33 µL deionized formamide) to prepare a 10 nM polyA RNA smFISH probe hybridization suspension, taking care to protect it from light. The dissected worms were washed with smFISH wash buffer (2x SSC, 10% formamide, 0.1% Tween-20) for 5 minutes at room temperature on a rocker. After removing excess buffer, 100 µL of the 10 nM polyA RNA smFISH probe suspension was added to each sample, thoroughly mixed, and the samples were incubated for 24 to 36 hours at 37 °C, with shaking at 850 rpm in the dark. After completing the probe hybridization, samples were washed once with 1 mL of smFISH wash buffer for 5 minutes and washed once with 1 mL of 1 μg/mL DAPI in smFISH wash buffer for 30 minutes in the dark. Then the samples underwent two additional washes with 1 mL of smFISH wash buffer for 5 minutes each. Excess buffer was carefully removed from the fixed samples. Samples were resuspended in 15 µL of ProLong Gold Antifade Mountant (Invitrogen P36934) and allowed to sit for 30 minutes. The resuspended gonads were then pipetted onto a microscope slide and covered with a 24 x 24 mm coverslip. Any excess liquid was removed carefully. Nail polish was applied along the edges of the coverslip to prevent drying and movement of the samples during imaging. The slides were cured at room temperature in the dark for approximately 24 hours before imaging.

## ACKNOWLEDGEMENTS

Some strains used in this study were provided by the CGC, which is funded by NIH Office of Research Infrastructure Programs (P40 OD010440). This work is supported in part by NIH NIH grant R01-GM132457 to H.-C.L.; the National Natural Science Foundation of China (grants 31922019) and the program for HUST Academic Frontier Youth Team (grant 2018QYTD11) to DZ.

## AUTHOR CONTRIBUTIONS

Conceptualization: D.Z., and H-C. L., Investigation: K.S., Y.Z., Z.D., and D.Z., Data Analyses: D.Z., K.S., Bioinformatics analyses: D.Z., Writing: H-C. L. and D.Z., Supervision: D.Z., and H-C. L., Funding acquisition: D.Z., and H-C. L.

## DATA AND CODE AVAILABILITY

The proteomic data are deposited to Proteomics Identification database (PRIDE) with the project accession number access number of PXD051572. The high-throughput sequencing data have been deposited to CNCB via the GSA data repository with accession number CRA016032 and can be found at https://ngdc.cncb.ac.cn/gsa/s/2HHgd5Ta. All custom scripts used for bioinformatic analysis are available at the supplemental data file. Any additional information required to reanalyze the data reported in this paper is available from the lead contact upon request.

## FIGURE LEGENDS

**Figure S1.**
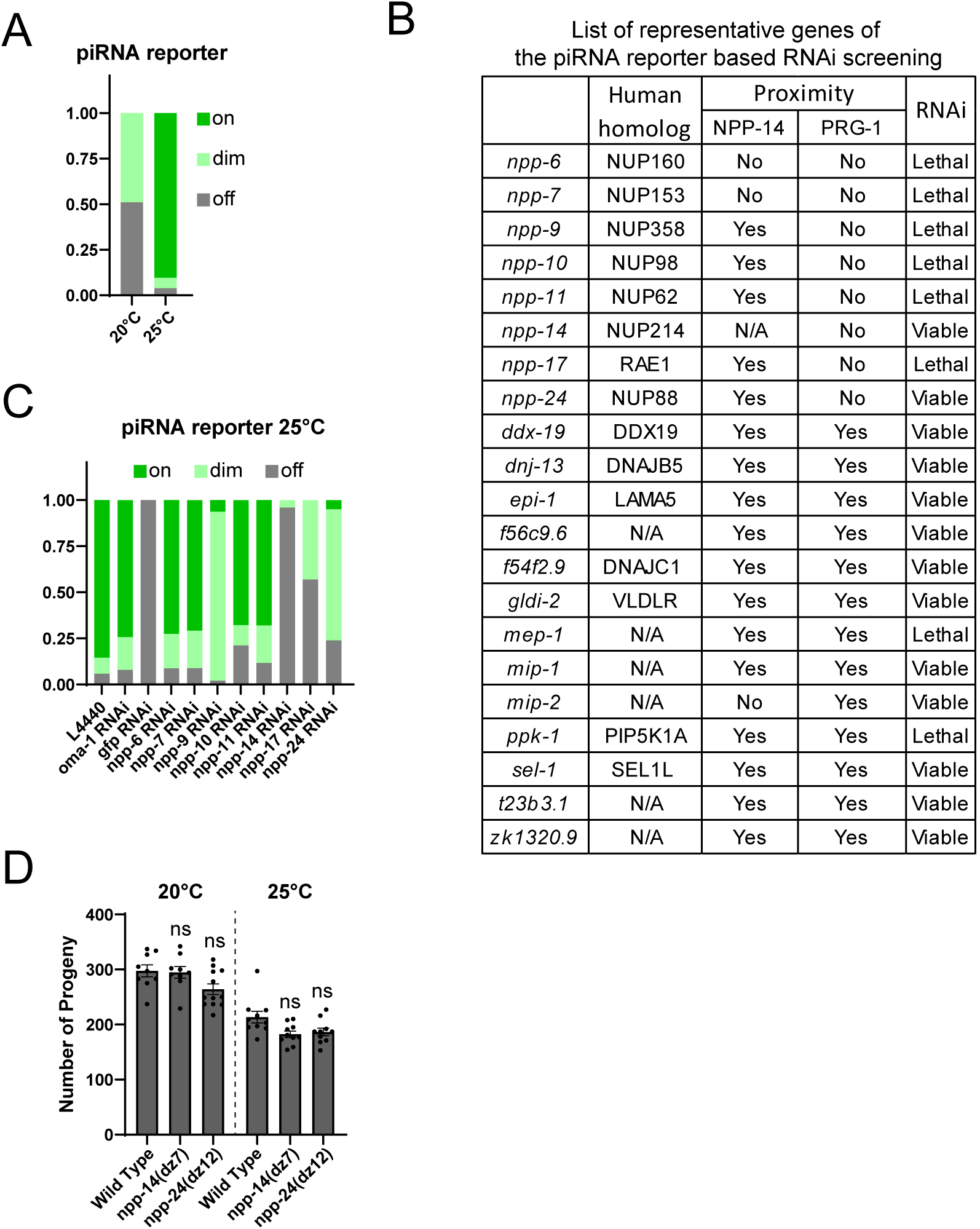
A candidate screen to examine factors that regulate piRNA silencing, related to Figure 1. (A) Percentage of screened animals displaying the indicated GFP expression levels of the piRNA reporter for worms grown at the indicated temperature. (B) List of candidate genes screen for the capability of piRNA reporter silencing at 25°C. Proximity labeling results of TurboID::NPP-14 and TurboID::PRG-1 are summarized. (C) Percentage of screened animals displaying the indicated GFP expression levels of the piRNA reporter in the indicated RNAi strains grown at 25°C. L4440 empty vector and RNAi against an unrelated gene, *oma-1*, were utilized as negative controls. RNAi against *gfp* served as the positive control. (D) Brood sizes of indicated strains cultured at 20°C or 25°C. One-way ANOVA followed by Dunnett’s correction for multiple comparisons was conducted. n=10.

**Figure S2.**
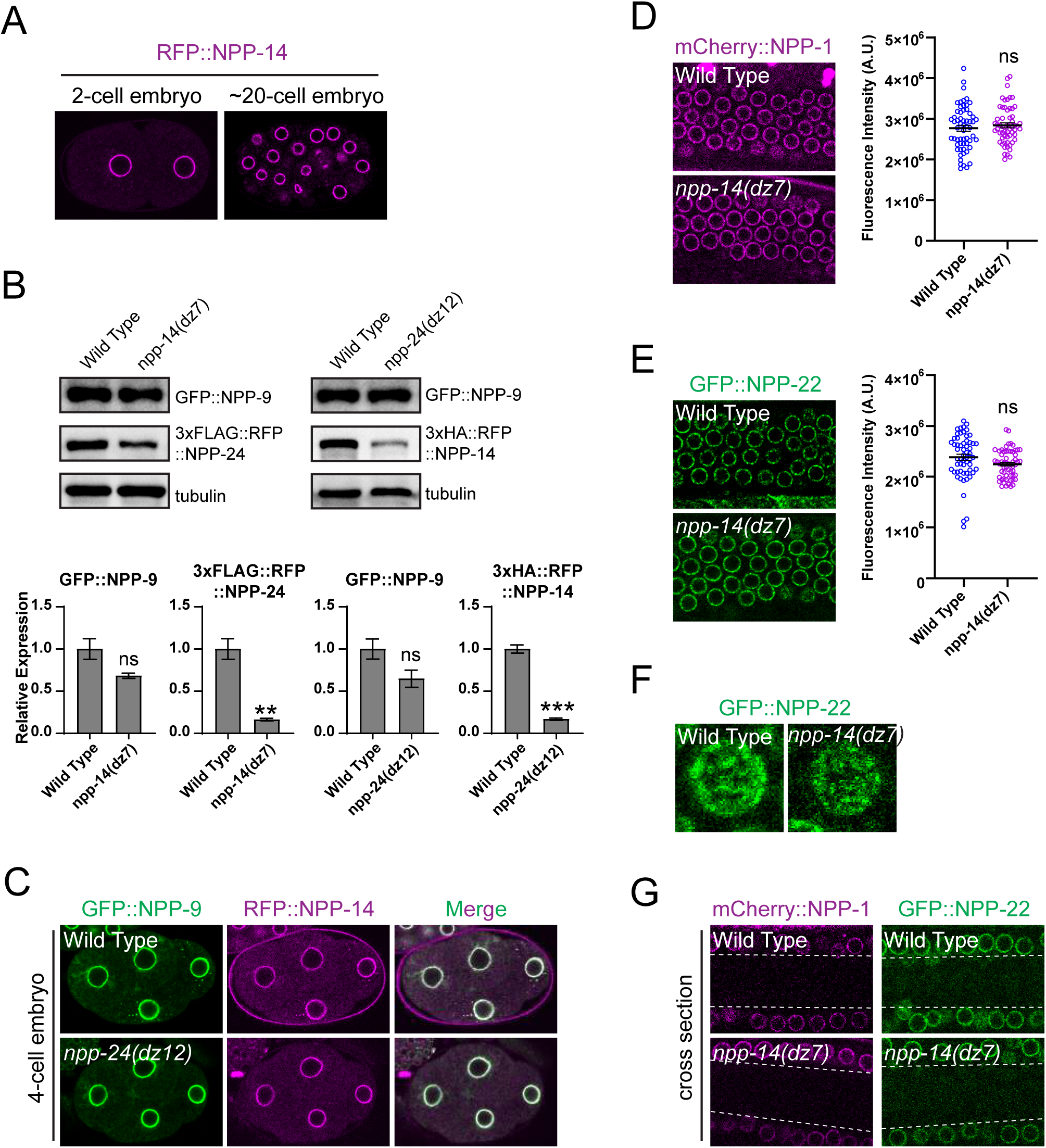
NPP-11 and NPP-24 promote NPC clustering and nuclear tethering in the germline, related to Figure 2. (A) Fluorescent micrographs showing the localization of RFP::NPP-14 in the 2-cell embryo and a the 20-cell embryo. (B) Western blot analyses showing the expression levels of the indicated proteins in the specified strains. Antibodies specifically against GFP, FLAG, HA, and tubulin were employed. Statistical analysis was conducted using a one-tailed Student’s t test by three independent blots. **, p<0.01; ***, p<0.001. (C) Fluorescent micrographs displaying the localization of RFP::NPP-14 and GFP::NPP-9 in the 4-cell embryo in the indicated strains. Note that RFP::NPP-14 remains stable in the 4-cell embryo in the *npp-24(dz12)* mutant. (D-E) Fluorescent micrographs showing the localization of (D) mCherry::NPP-1 and (E) GFP::NPP-22 in the indicated strains. Mean fluorescent intensity of granules is analyzed using a one-tailed Student’s t-test. n=12-15. Enrichment of mCherry::NPP-1 and GFP::NPP-22 is not altered by depletion of NPP-14. (F) Fluorescent micrographs showing the clustering of GFP::NPP-22 in the indicated strains. The surface sections of germline nuclei from the pachytene region are shown. (G) Fluorescent micrographs of the germline cross-section showing mCherry::NPP-1 and GFP::NPP-22 in the indicated strains. The area between two dashed lines is germline syncytial cytoplasm.

**Figure S3.**
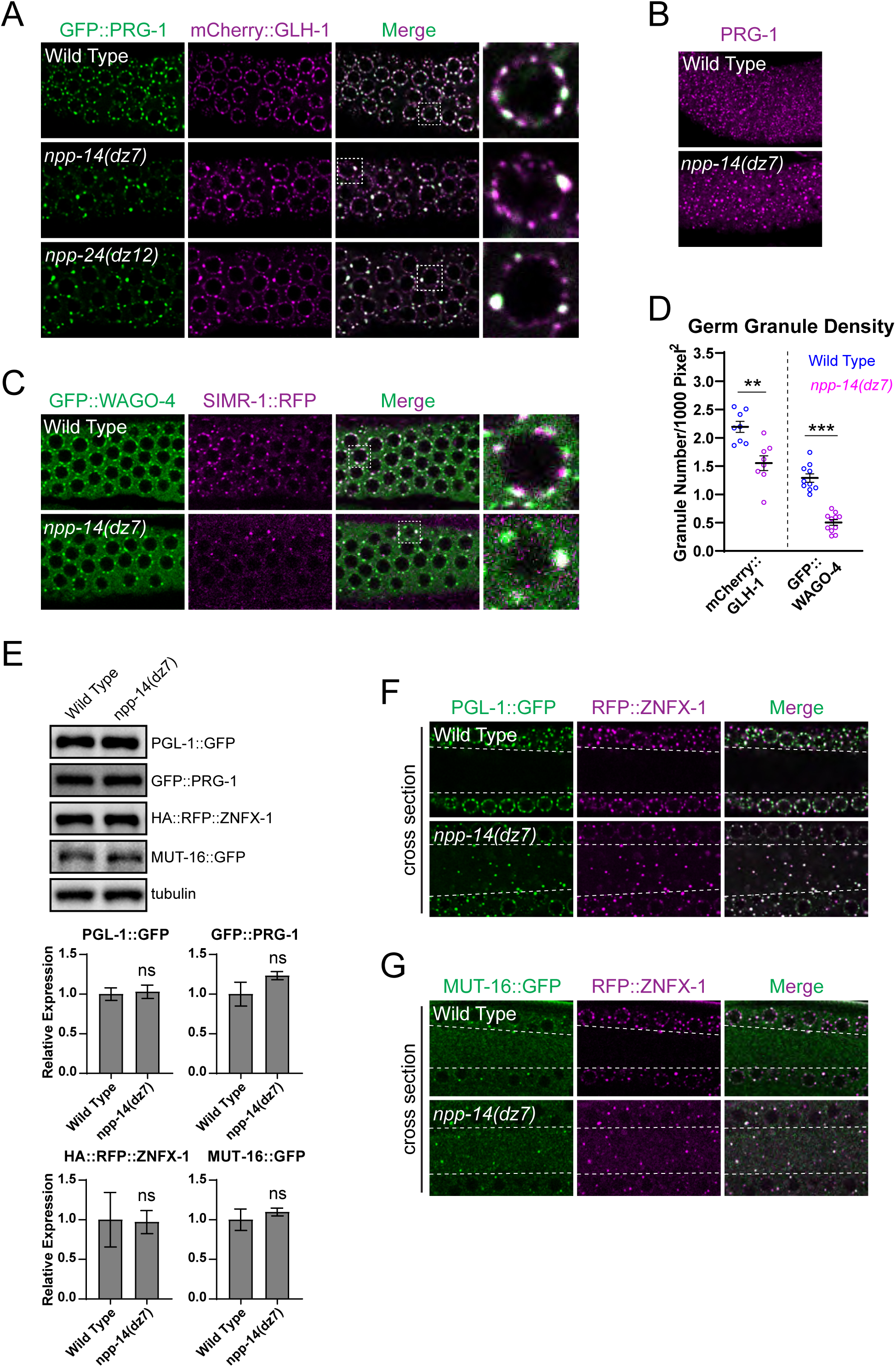
Diverse roles of NPP-14 and NPP-24 in germ granule formation, related to figure 3. (A) Fluorescent micrographs showing the localization of GFP::PRG-1 and mCherry::GLH-1 in the indicated strains. (B) (Immunofluorescence micrographs of endogenous PRG-1 in the indicated strains. Antibodies against PRG-1 were employed. (C) Fluorescent micrographs showing the colocalization of GFP::WAGO-4 and SIMR1::RFP in the indicated strains. (D) Mean granule density in the indicated strains. Statistical analysis was conducted using a one-tailed Student’s t-test. n=12-15. **, p<0.01; ***, p<0.001. (E) Western blot analyses showing the expression levels of the indicated proteins in the specified strains. Antibodies against GFP, HA, and tubulin were employed. Statistical analysis was conducted using a one-tailed Student’s t test by three independent blots. **, p<0.01; ***, p<0.001. (F-G) Fluorescent micrographs of the germline cross-section showing the localization of the specified proteins in the indicated strains. The area between two dashed lines is germline syncytial cytoplasm.

**Figure S4.**
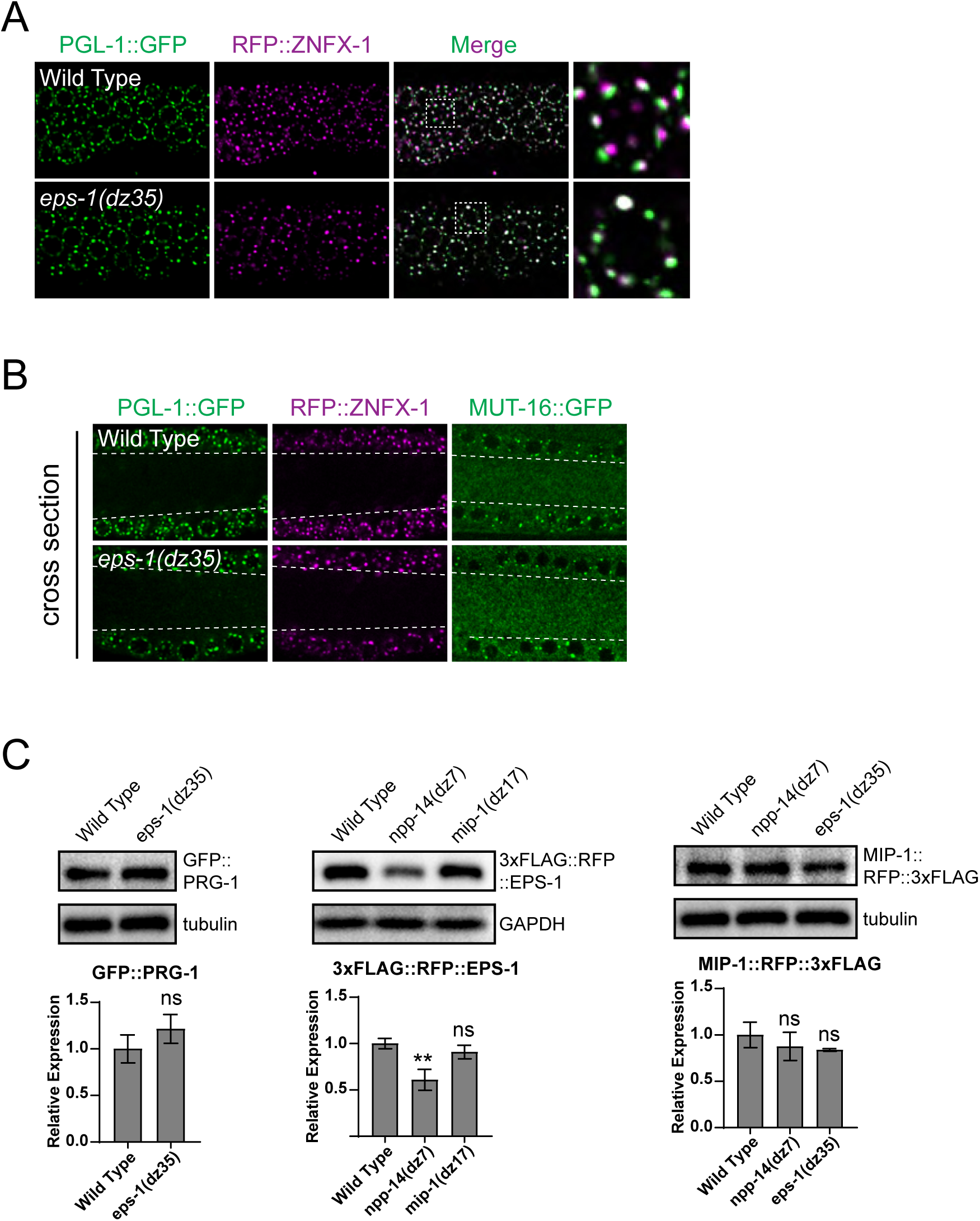
The roles of EPS-1 in germ granule formation and organization, related to Figure 5. (A) Fluorescent micrographs showing the localization of PGL-1::GFP and RFP::ZNFX-1 in the indicated strains. (B) Fluorescent micrographs of the germline cross-section showing the localization of PGL-1::GFP and RFP::ZNFX-1 in the indicated strains. The area between two dashed lines is germline syncytial cytoplasm. (C) Western blot analyses showing the expression levels of the indicated proteins in the specified strains. Antibodies against GFP, FLAG, tubulin, and GAPDH were employed. Statistical analysis was conducted using a one-tailed Student’s t-test or one-way ANOVA followed by Dunnett’s correction for multiple comparisons by three independent blots. **, p<0.01.

**Figure S5.**
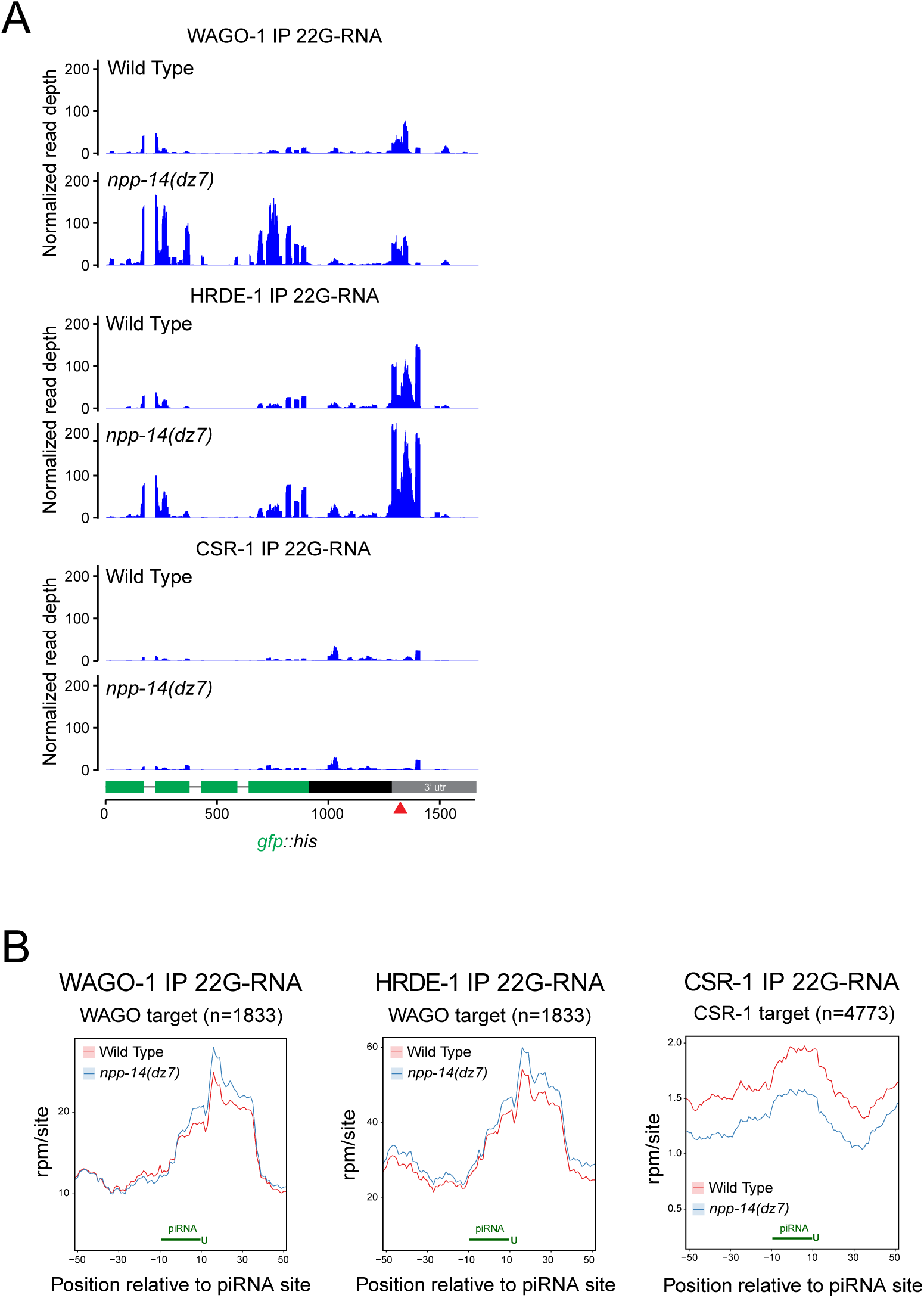
WAGO-associated 22G-RNAs are over-amplified at the piRNA targeting sites in the *npp-14* mutant, related to Figure 6. (A) Distribution of WAGO-1 (top), HRDE-1 (middle), or CSR-1 (bottom)-bound 22G-RNAs mapped to the piRNA reporter sequences in the indicated strains. The red arrowhead indicates the location of sequences complementary to the endogenous piRNA 21ur-1. (B) Density of WAGO-1 (left)-, HRDE-1 (middle)- or CSR-1 (right)-bound 22G-RNAs within a 100-nt window around predicted piRNA targeting sites of the specified Argonaute target genes in the indicated strains.

**Figure S6.**
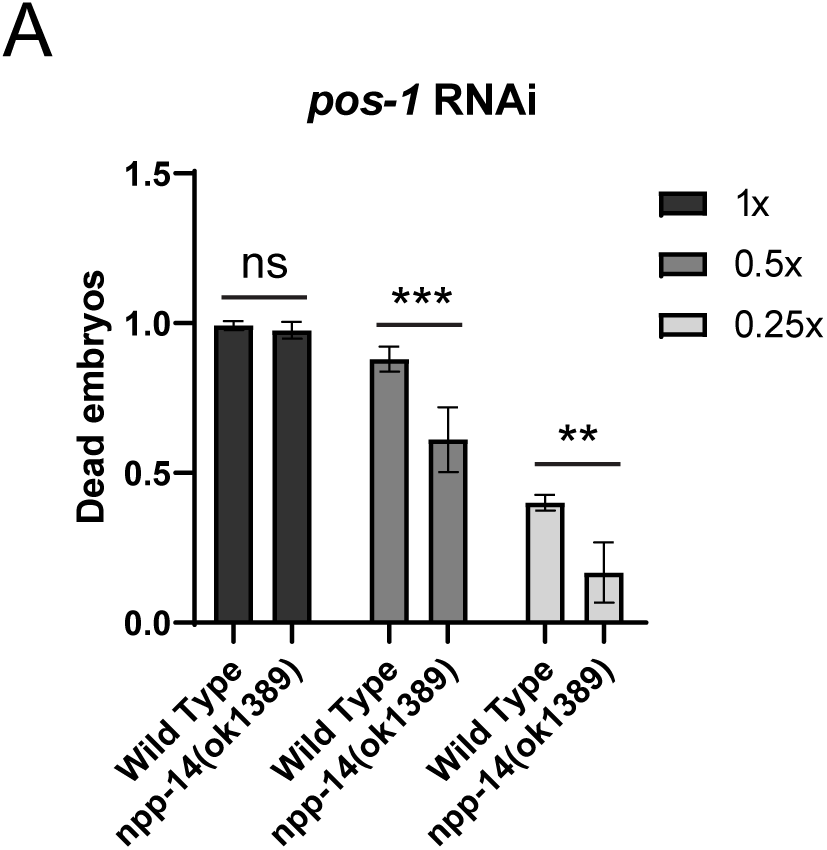
RNAi is compromised in the *npp-14* mutant, related to Figure 7. (A) Serial dilutions of bacteria expressing the *pos-1* dsRNA are employed for the RNAi feeding treatments in the indicated strains.

## SUPPLEMENTARY TABLES

**Supplementary Table 1. List of proteins identified in NPP-14 and PRG-1 complex.**

**Supplementary Table 2. List of C. elegans strains used in this study. Supplementary Table 3. List of primers and oligos used in this study.**

